# Toxic anti-phage defense proteins inhibited by intragenic antitoxin proteins

**DOI:** 10.1101/2023.05.02.539157

**Authors:** Aoshu Zhong, Xiaofang Jiang, Alison B. Hickman, Katherine Klier, Gabriella I. C. Teodoro, Fred Dyda, Michael T. Laub, Gisela Storz

**Author notes:** Current address: Freshwater and Marine Sciences, University of Wisconsin-Madison, Madison, WI, USA.

## Abstract

Recombination-promoting nuclease (Rpn) proteins are broadly distributed across bacterial phyla, yet their functions remain unclear. Here we report these proteins are new toxin-antitoxin systems, comprised of genes-within-genes, that combat phage infection. We show the small, highly variable Rpn *C*-terminal domains (Rpn_S_), which are translated separately from the full-length proteins (Rpn_L_), directly block the activities of the toxic full-length proteins. The crystal structure of RpnA_S_ revealed a dimerization interface encompassing a helix that can have four amino acid repeats whose number varies widely among strains of the same species. Consistent with strong selection for the variation, we document plasmid-encoded RpnP2_L_ protects *Escherichia coli* against certain phages. We propose many more intragenic-encoded proteins that serve regulatory roles remain to be discovered in all organisms.

**Significance:** Here we document the function of small genes-within-genes, showing they encode antitoxin proteins that block the functions of the toxic DNA endonuclease proteins encoded by the longer *rpn* genes. Intriguingly, a sequence present in both long and short protein shows extensive variation in the number of four amino acid repeats. Consistent with a strong selection for the variation, we provide evidence that the Rpn proteins represent a phage defense system.

---

The annotation of bacterial genes generally assumes that each coding sequence directs the synthesis of one protein product. However, there are a few examples where two protein products are translated from the same gene, one product corresponding to the full open reading frame (ORF) and the second translated from an internal translation initiation start (iTIS) (1). The functions of most products of these genes-within-genes are not known, though for the few that have been characterized, the two products can have complementary or opposing functions. One example of a complementary function is provided by a *Synechocystis* type I-D CRISPR-Cas Cascade system, where the full-length Cas10d protein forms a complex with Cas11d, which is translated from an iTIS (2). The complex is required for specific DNA binding by the type I-D Cascade complex, and without Cas11d, the Cascade complex has little or no DNA binding activity. Recent experiments to examine transcriptome-wide ribosome binding (ribo-seq) in the presence of inhibitors that trap the ribosome on translation initiation sites suggested there are more iTIS than initially considered (3, 4). The five *rpn* (recombination-promoting nuclease) genes of *E. coli* each have an iTIS that could potentially direct the synthesis of small proteins corresponding to the variable *C*-terminal tail (Fig. 1*A* and *SI Appendix*, Fig. S1*A*).

**Fig. 1.**
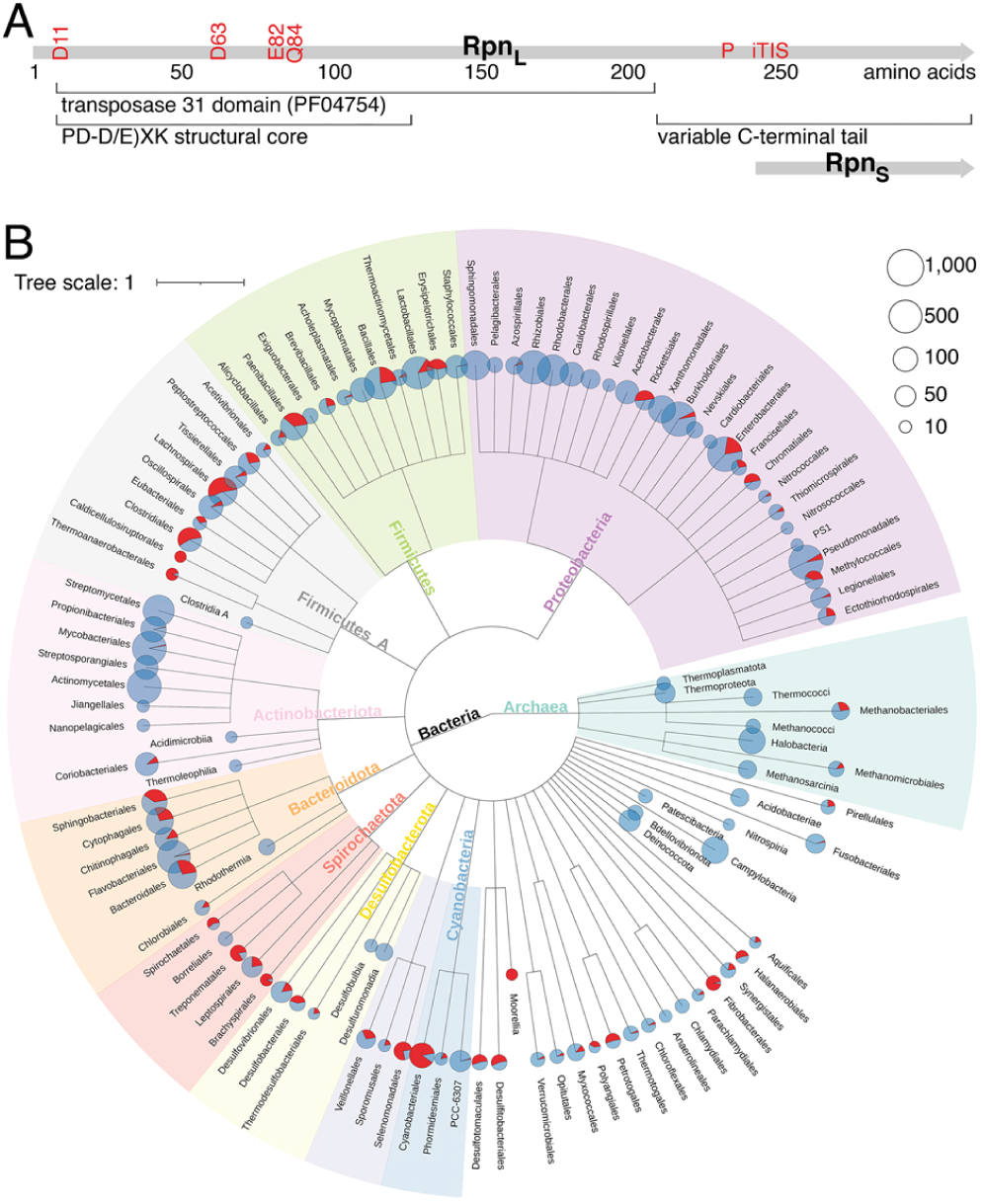
Broadly-distributed *rpn* genes move by horizontal gene transfer. (*A*) Diagram of *rpn* genes adapted from (7). Positions of catalytic residues for RpnA and internal promoter (P) and iTIS are indicated in red font. (*B*) Species distribution of *rpn* orthologs. Clades with more than 15 children are displayed. The pie charts represent the percentage of species with (red) and without (blue) a *rpn* ortholog. The size of the pie chart represents the number of species with *rpn* orthologs in that clade.

Rpn proteins, which are members of the diverse PD-(D/E)XK phosphodiesterase superfamily (5), have been proposed to be involved in horizontal gene transfer based on the report that the conserved *N*-terminal domain has homology to transposases (6). Previous studies showed that overexpression of *E. coli* Rpn proteins reduced cell viability in a *recA*^-^ background and induced the DNA damage (SOS) response in a *recA*^+^ host (7). Furthermore, the purified RpnA protein was shown to possess DNA endonuclease activity. These observations support the idea that Rpn proteins are DNA-mobilizing enzymes, but the physiological role of this DNA cleavage activity is unknown. Thus, we set out to characterize the long, full-length proteins (Rpn_L_) as well as the small *C*-terminal proteins (Rpn_S_), which we hypothesized are translated from the iTIS and could act in conjunction with the Rpn_L_ proteins.

## Results

### *rpn* Genes Move by Horizontal Gene Transfer but Do Not Encode Conventional Transposases

Phylogenetic analysis revealed that members of the *rpn* orthologous gene cluster (COG5464) are widely distributed and are found in 34 bacterial and two archaeal phyla (Fig. 1*B*). Their distribution is strongly non-uniform indicating that *rpn* genes are prone to frequent horizontal gene transfer. Additionally, the number of *rpn* genes can vary between different strains of the same species. For example, no *rpn* genes were found in the genome of *Bacillus thuringiensis serovar kurstaki str. T03a001* (assembly accession: GCF_000161575.1) while 26 copies are present in *Bacillus thuringiensis serovar yunnanensis* (assembly accession: GCF_002147825.1).

*rpn* genes are commonly annotated as encoding a “putative transposase”, and genomic synteny among strains differing in *rpn* genes suggests the *rpn* genes were recently acquired or lost as a single gene cluster, a pattern similar to insertion sequences (8). However, unlike typical insertion sequences, the coding regions of *rpn* genes are not flanked by identifiable inverted terminal repeats or target-site duplications. Furthermore, comparative genomic analysis of closely related genomes with and without *rpn* genes shows that the ends of the genomic region containing the *rpn* gene vary, contrasting with insertion sequences where the ends are generally defined. In genomes where *rpn* gene arrays were found, the arrays appear to be generated by tandem direct duplications of individual genomic regions rather than by transposition of different *rpn* genes into the same genomic locus given the absence of target-site duplications and the presence of different types of duplications (*SI Appendix*, Fig. S1*B*). Although the TnsA protein of the heteromeric Tn7 transposase has a PD-(D/E)XK catalytic domain, there are no known autonomous transposases with this protein fold (9, 10), and we find no evidence that *rpn* genes are capable of autonomous transposition. We therefore think it is highly unlikely that Rpn proteins function as transposases.

### Small Proteins Are Translated from Within *rpn* Genes

We next wanted to determine the role of the iTIS (Fig. 2*A*) and ribosome profiling signal detected for all five *E. coli rpn* genes after treatment with the translation inhibitors Onc112 or retapamulin (3, 4) (Fig. 2*B*, *D*, *F*, and *G* and *SI Appendix*, Fig. S2*A*). To see if small proteins corresponding to the *C*-terminal tails are synthesized, we introduced a sequential peptide affinity (SPA) tag on the chromosome upstream of each *rpn* stop codon, permitting detection via immunoblot analysis. In cells grown to exponential (“exp”) or stationary (“stat”) phase, all five small proteins corresponding to the *C*-terminus (denoted Rpn_S_) were observed at different levels (Fig. 2*C*, *E*, and *H* and *SI Appendix*, Fig. S2*B*). Among the full-length proteins (denoted Rpn_L_), only RpnB_L_, RpnC_L_ and RpnE_L_ were detected under the growth conditions examined (Fig. 2*E* and *H*). The higher levels of the Rpn_S_ proteins are consistent with previous differential RNA-seq (dRNA-seq) data (11), which shows a transcription start site within all *rpn* genes for a short transcript expressed at higher levels than the full-length mRNA. The introduction of mutations in the ribosome binding site (RBS) and predicted iTIS with minimal change to the *rpnA* coding sequence (RpnA_K240R+E241D+M244L_, hereafter denoted RpnA_L*\**_) on the *E. coli* chromosome (Fig. 2*A*) abolished RpnA_S_ expression (Fig. 2*C*), confirming the RpnA_S_ protein is translated from the predicted iTIS.

**Fig. 2.**
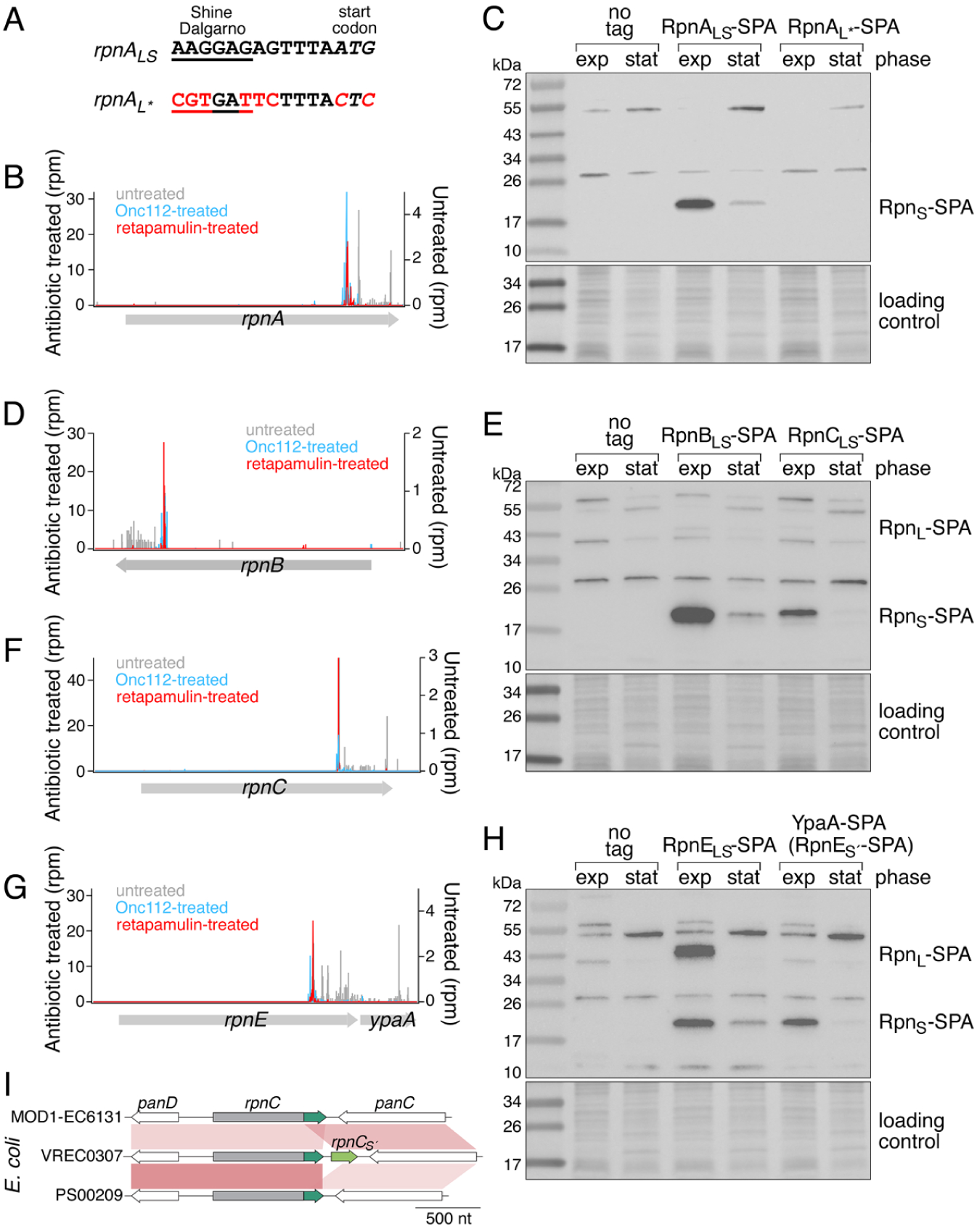
The *rpn* genes encode smaller proteins (Rpn_S_). (*A*) Sequence of RpnA_S_ ribosome binding site, and mutations to eliminate *rpnA_S_* ribosome binding and start codon. Browser images of ribosome profiling data for *rpnA* (*B*), *rpnB* (*D*), *rpnC* (*F*), *rpnE*-*ypaA/rpnE_Ś_* (*G*). Ribosome density for an untreated control (gray) and cells treated with Onc112 (blue) (4) or retapamulin (red) (3) are shown. **c**, **e**, **h**, Immunoblot analysis of the levels of SPA-tagged RpnA_S_ (*C*) and RpnB_S_, and RpnC_S_ (*E*) RpnE_S_ and YpaA/RpnE_Ś_ (*H*). *E. coli* MG1655 strains were grown to exponential (exp) and stationary (stat) phase in LB. The SPA tag was detected with monoclonal anti-FLAG M2-peroxidase (HRP) antibody. Ponceau S staining of membranes served as loading controls. (*I*) Diagram showing an example of insertion and deletion of *rpn_Ś_* genes (green) among strains of *E. coli*, with red connections linking orthologous regions and color indicating identity (darker=higher, lighter=lower).

We also wanted to determine whether an Rpn_S_ protein is expressed from *rpn* genes from other organisms and thus cloned an *rpn* gene from the Gram^+^ bacterium *Clostridioides difficile* into a plasmid that we introduced into *E. coli*. For this construct, we also observed expression of an Rpn_S_ protein, which again was eliminated by the introduction of mutations of the predicted RBS and iTIS (*SI Appendix*, Fig. S2*C* and *D*). This observation suggests co-expression of Rpn_L_ and Rpn_S_ proteins is broadly conserved.

### Varied Expression of Small Proteins

In an analysis of the *rpn* genes present in diverse *E. coli* strains (*SI Appendix*, Fig. S3*A* and *B*), we observed that while most of the genes are encoded on the chromosome, some are encoded on plasmids. Intriguingly, on both the chromosome and plasmids, the Rpn_S_ protein can be expressed from either within an *rpn* gene or as a separate gene, annotated as encoding proteins with the DUF4351 domain of unknown function (PF14261) in some species (Dataset S1). For example, we noted that a second copy of RpnE_S_ was encoded by the *ypaA* gene (denoted *rpnE_Ś_* here) downstream of the *rpnE* gene. RpnE_Ś_ was detected upon SPA tagging indicating the protein is synthesized (Fig. 2*H*). A second copy of *rpnC_S_* is similarly found downstream of *rpnC* in some *E. coli* strains (Fig. 2*I*). Phylogenetic analysis, as demonstrated in *SI Appendix*, Fig. S3*C*, revealed that the downstream small protein homologs RpnC_Ś_ and RpnE_Ś_ cluster together with their respective upstream small protein homologs, RpnC_S_ and RpnE_S_, rather than clustering with each other. This observation suggests that the origin of the downstream small proteins occurred independently, with RpnC_Ś_ originating from RpnC and RpnE_Ś_ originating from RpnE. The *rpn_S_*’ gene also can be lost through the deletion of the encoding region or fusion of the *rpn N*-terminal domain and *rpn_S_*’ encoding region (Fig. 2*I*).

For further analysis of Rpn protein function, we expressed both RpnA and RpnB proteins from a rhamnose-inducible promoter as done previously (here denoted as expressing RpnA_LS_ or RpnB_LS_) (7) or from their own promoters on a low copy plasmid. We also expressed plasmid-encoded RpnP2_LS_ from its own native promoter on a low copy plasmid. To first compare the relative levels of the proteins expressed from these constructs, we again integrated an SPA tag upstream of the stop codon. Intriguingly, the relative levels of Rpn_L_ and Rpn_S_ varied widely with higher levels of Rpn_L_ than Rpn_S_ from the rhamnose-inducible promoter than the native promoter, except for RpnP2_L_ (*SI Appendix*, Fig. S2*E* and *F*). The reasons for the differential expression are not known but suggest additional regulation.

### Rpn_S_ Proteins Block Rpn_L_-Dependent Growth Inhibition

Although previous studies showed that Rpn proteins expressed from the rhamnose promoter reduce cell viability when overexpressed (7), Rpn_S_ likely was unknowingly co-expressed from these constructs, complicating the interpretation of these results. To deconvolute the effects of the two proteins, we constructed plasmids carrying *rpnA* or *rpnB* with mutated iTIS and RBS sequences (RpnA_L*_ or RpnB_L*_) to fully abolish the expression of RpnA_S_ or RpnB_S_. When assessed by cell density measurements in liquid culture, there was no effect on *E. coli* ER2170 growth (Fig. 3*A*) with the empty plasmid (black) or RpnA_LS_ overexpression (green), while a defect was observed when expressing RpnA_L*_ (red). For the RpnB constructs (Fig. 3*B*), we observed some growth defect for cells expressing RpnB_LS_ (green), but the effect was even stronger for overexpression of RpnB_L*_ alone (red). For RpnP2 (Fig. 3*C*), all constructs for RpnP2_L*_ expressed from its native promoter had additional inactivating site mutations indicating strong selection against expression of RpnP2_L_ alone. We were only able to obtain the intact RpnP2_L*_ construct when the *rpnP2* gene was cloned behind the P_BAD_ promoter, which is repressed by glucose and activated by arabinose. In the presence of glucose, the RpnP2_L*_ cells showed a slight growth defect (top panel, red). In contrast, growth was strongly inhibited by RpnP2_L*_ upon the addition of arabinose (bottom panel, red), while RpnP2_LS_ cells (green), which also expressed the RpnP2_S_ protein, grew like vector control cells (black) under both conditions. Thus, Rpn proteins have the properties of toxin-antitoxin systems (12) with Rpn_L_ proteins inhibiting growth and Rpn_S_ proteins serving as the cognate antitoxin.

**Fig. 3.**
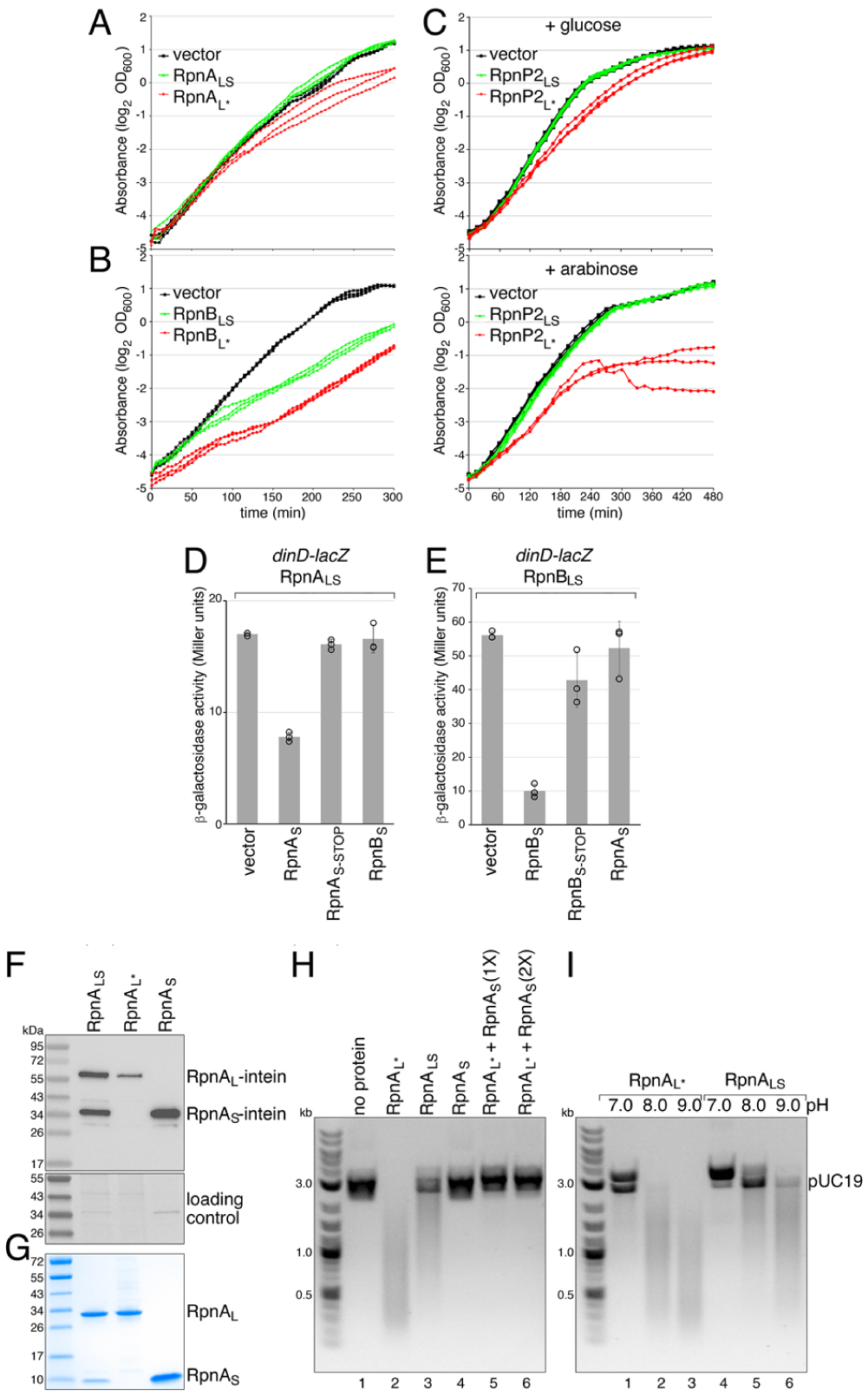
Rpn_S_ proteins function as antitoxins. RpnA_L_ (*A*) and RpnB_L_ (*B*) inhibit *E. coli* growth. *E. coli* ER2170 cells harboring indicated plasmids were grown in LB. Three biological replicates are shown. (*C*) RpnP2_L_ inhibits *E. coli* growth. *E. coli* MG1655 cells harboring indicated plasmids were grown in LB with added glucose or arabinose. For *A*, *B*, and *C*, three biological replicates are shown. (*D*) RpnA_S_, but not RpnB_S_, blocks RpnA_L_ induction, and (*E*) RpnB_S_, but not RpnA_S_, blocks RpnB_L_ induction of the *dinD-lacZ* reporter of the SOS DNA damage response. RpnA_LS_ and RpnB_LS_ are overexpressed from the rhamnose-inducible P*_rhaB_* promoter. RpnA_S_ and RpnB_S_ are overexpressed from the arabinose-inducible P_BAD_ promoter. For (*D*) and (*E*), average of three independent biological repeats is given with standard deviation. For (*A*)-(*E*), the ER2170 or MG1655 strain backgrounds were WT for the *rpn* genes. (*F*) Immunoblot analysis of intein-tagged RpnA_LS_, RpnA_L*_ and RpnA_S_ overexpression. Ponceau S staining of membrane served as loading control. (*G*) Coomassie-stained Tris-glycine SDS-PAGE gel of purified RpnA_LS_, RpnA_L*_ and RpnA_S_. (*H*) Purified RpnA_S_ blocks dsDNA endonuclease activity of RpnA_L*_. Indicated purified proteins were incubated with pUC19. (*I*) RpnA_L_ activity is pH-dependent. Purified RpnA_L*_ (lanes 1-3) or RpnA_LS_ (lanes 4-6) were incubated with pUC19 at pH 7.0, pH 8.0 or pH 9.0. Images in (*H*) and (*I*) are inverted from the original. Each of the nuclease assays was repeated at least twice.

### Rpn_S_ Proteins Block Rpn_L_-Dependent SOS Induction

To test whether the Rpn_S_ proteins can block the DNA damage caused by Rpn_LS_ overexpression reported previously (7), we employed a two-plasmid system to separately overexpress Rpn_S_. We observed that, when the proteins are overexpressed, RpnA_LS_ and RpnB_LS_ both induce the SOS response (*SI Appendix*, Fig. S4*A*), as monitored by the chromosomally-encoded, DNA damage-inducible P*_dinD_*-*lacZ* reporter (7). For these experiments, induction of RpnA_LS_ and RpnB_LS_ was titrated to allow for overexpression without significantly impacting growth. Co-expression of RpnA_S_ or RpnB_S_ reduced SOS induction in the corresponding Rpn_LS_ strain (Fig. 3*D* and *E*). This block is no longer observed when a stop codon was introduced in the RpnA_S_ (at E25) or RpnB_S_ (at D29) coding sequence indicating that the repressive effect is due to the protein and not due to overexpression of the RNA. The repressive effect of each small protein is specific to the Rpn_L_ protein encoded by the same sequence as overexpression of RpnA_S_ did not block RpnB_LS_-dependent *dinD-lacZ* induction while RpnB_S_ did not block RpnA_LS_-dependent induction. These observations further support that RpnA_S_ and RpnB_S_ specifically block the activities of the corresponding larger protein.

### RpnA_S_ Blocks RpnA_L_-Dependent DNA Cleavage In Vitro

To directly examine the effect of the Rpn_L_ and Rpn_S_ proteins on DNA cleavage, RpnA derivatives, which were easier to express given that they were the least toxic, were overexpressed as *C*-terminally tagged intein fusion proteins and purified. When the WT *rpnA* gene was cloned into an overexpression vector, the *C*-terminally tagged RpnA_L_ and RpnA_S_ (61 kDa and 33 kDa, respectively) were both detected (Fig. 3*F*) and purified together (Fig. 3*G*). Thus, we also overexpressed and purified the RpnA_L*_ and RpnA_S_ proteins separately (Fig. 3*F* and *G*). In assays for DNA cleavage activity using both double-stranded (Fig. 3*H*) and single-stranded (*SI Appendix*, Fig. S4*B*) DNA substrates, the RpnA_L*_ protein had strong DNA cleavage activity (lane 2). Consistent with the conclusion that RpnA_S_ blocks RpnA_L_ activity, we found that RpnA_LS_ has significantly less endonuclease activity (lane 3), and the addition of RpnA_S_ to RpnA_L*_ completely blocks the cleavage (lanes 5 and 6). Interestingly, the DNA cleavage activities of both RpnA_L*_ and RpnA_LS_ are higher at a more alkaline pH (Fig. 3*I*) as was previously observed for RpnA_LS_ (7), suggestive of a role for the cleavage activity at higher pH or other specific growth condition with similar consequences.

### RpnA_S_ Forms a Complex with RpnA_L_

To examine whether RpnA_S_ is inhibiting RpnA_L_ through a direct interaction, the oligomerization states of the purified proteins were assessed by size exclusion chromatography (Fig. *4A*). The individually purified RpnA_L*_ (33.3 kDa) and RpnA_S_ (5.4 kDa) proteins eluted as distinct peaks. However, when RpnA_L*_ and RpnA_S_ were mixed, a stable RpnA_L*_-RpnA_S_ complex was observed that eluted at the same volume (arrow) as the native RpnA_LS_ complex. SDS-PAGE analysis of the peak fractions confirmed that both RpnA_L*_ and RpnA_S_ are in the corresponding fractions (*SI Appendix*, Fig. S5*A*). In a multi-angle light scattering coupled with size exclusion chromatography (SEC-MALS) experiment, the molecular weight of RpnA_S_ was determined to be 10.2 ± 0.2 kDa, consistent with a dimer. The molecular weight of the RpnA_LS_ complex was determined to be 74.1 ± 0.5 kDa by SEC-MALS, consistent with a tetramer of two RpnA_S_ subunits and two RpnA_L_ subunits. Intriguingly, the RpnA_LS_ complex is stable at pH 7.0 and 8.0, yet the subunits dissociate at pH 9.0 and 10.0 (*SI Appendix*, Fig. S5*B*), paralleling the increase in DNA cleavage activity observed for RpnA_LS_ (Fig. 3*I*).

**Fig. 4.**
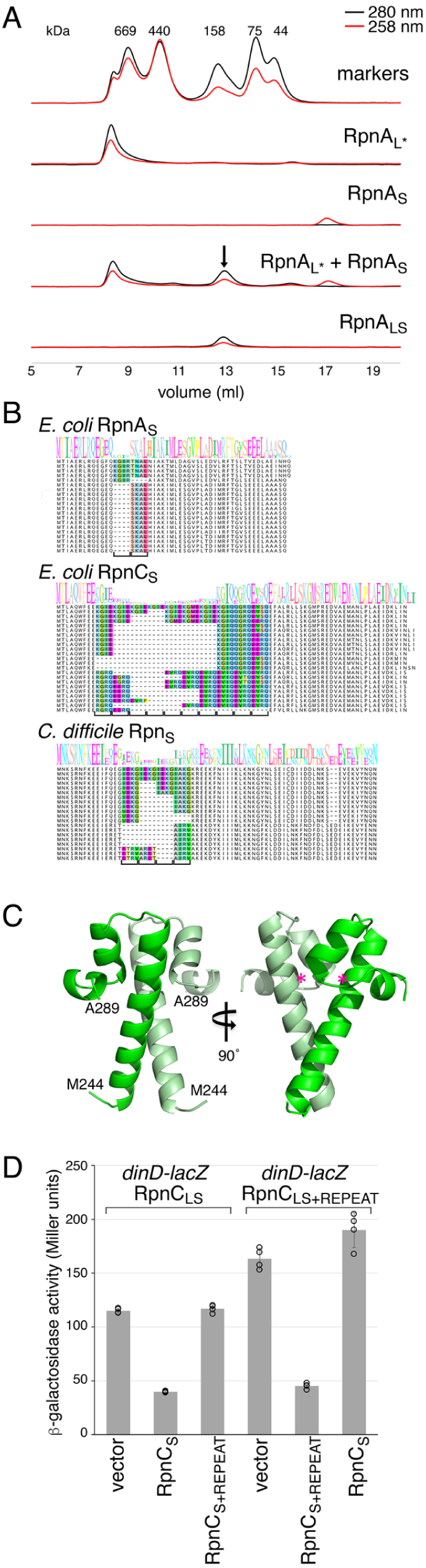
RpnA_L_ and RpnA_S_ form a complex. (*A*) Size exclusion chromatograms of indicated proteins on a Superdex 200 column. Although the RpnA_LS_ peak migrates similarly to the 158 kDa marker peak, SEC-MALS analysis, which is less influenced by protein shape, indicates that the complex has a molecular weight of 74.1 ± 0.5 kDa. The RpnA_L*_ protein migrates as a large molecular weight complex indicative of an aggregate, but we do not observe precipitates of this protein. (*B*) Sequence alignments for *E. coli* RpnA_S_ and RpnC_S_ as well as *C. difficile* Rpn_S_. The sequence logo was built based on all the sequences for each group though only representative individual sequences are shown. The four amino acid repeat that varies between strains is colored in the alignment, with a bracket indicating the number of repeats. The alignments were manually curated. (*C*) Crystal structure of RpnA_S_. Pink asterisks represents approximate position where four amino acid insertions occur (H261 in RpnA_L_). (*D*) Only RpnC_S_ with same number of four amino acid repeats blocks RpnC_L_ induction of *dinD-lacZ*. RpnC_LS_ and RpnC_LS+REPEAT_ are expressed from the rhamnose-inducible P*_rhaB_* promoter, while RpnC_S_ and RpnC_S+REPEAT_ are expressed from the arabinose-inducible P_BAD_ promoter. For the REPEAT derivatives, a four amino acid repeat (KGIE) is inserted in the same position. Average of four independent biological repeats is given with standard deviation.

### *C*-terminal Domains Vary by Four Amino Acid Repeats and Comprise an Oligomerization Domain

A striking feature of the amino acid sequence present in both the Rpn_L_ and Rpn_S_ proteins, which is observed from the alignment of homologs from different strains of the same species, is variation by four amino acid repeats generally comprised of one hydrophobic amino acid and three charged residues or glycine (Fig. 4*B* and *SI Appendix*, Fig. S6). This is seen for all the *E. coli* Rpn_S_ proteins including Rpn_Ś_ proteins as well as those found in distantly related bacteria such as *Bacillus cereus*, *Bacteroides fragilis*, *C. difficile* and *Leptospira interrogans*.

Given the interaction between RpnA_L_ and RpnA_S_, we hypothesized that this hypervariable four amino acid repeat region present in both forms of the Rpn proteins might comprise an oligomerization domain. To gain insights into the domain, we solved the structure of RpnA_S_ at a resolution of 1.9 Å using X-ray crystallography (Fig. 4*C*, *SI Appendix*, Table S1). Consistent with the SEC-MALS data, RpnA_S_ is a dimer in the crystals in which the two monomers each fold as a compact three α-helix bundle. The long first helix of each monomer interacts with the other, burying a total surface area of 976 Å^2^. In all Rpn_S_ proteins where they have been inserted, the four amino acid repeats begin 8-20 residues after the initiating methionine (*SI Appendix*, Fig. S6), which places the insertion point for the hypervariable region within the first α-helix of Rpn_S_. Based on three-dimensional structure predictions by AlphaFold2 (13), the inserted repeats most likely increase the length of the α-helix rather than disrupt it (*SI Appendix*, Fig. S7).

To examine the consequences of changing the number of four amino acid repeats, we generated derivatives of the RpnC_LS_ and RpnC_S_ proteins, which have the highest number of repeats among *E. coli* strains, by adding one repeat. Interestingly, when assessed in the SOS response assay, the RpnC_S+REPEAT_ can no longer counteract RpnC_LS_ (Fig. 4*D*). Conversely, RpnC_S_ cannot counteract RpnC_LS+REPEAT_, but repression is restored when the two proteins with the extra repeat are expressed together. These observations are consistent with the repeat regions affecting the binding between the two Rpn_S_ monomers and providing specificity for the interaction between the Rpn_L_ and Rpn_S_ proteins.

### Rpn Proteins Contribute to Anti-phage Defense

The *rpn* genes share characteristics with known anti-phage defense systems such as restriction-modification enzymes and toxin-antitoxin complexes, including the ability to quickly diversify and extensive horizontal mobility. The genes also show a tendency to cluster with other defense genes (Fig. 5*A*), including systems with dCas12a or Type I-E Cascade (14). Like most anti-phage defense systems (15, 16), the *rpn* genes are not autonomously mobile but likely rely on other mechanisms such as recombination or “hitchhiking” with mobile elements to move. Given these similarities and the rapid change in the numbers of four amino acid repeats among Rpn_S_ proteins, which we surmised must be due to strong selective pressure acting on the *rpn* genes to diversify the *C*-terminal domain, we hypothesized Rpn proteins might defend against phage infection.

**Fig. 5.**
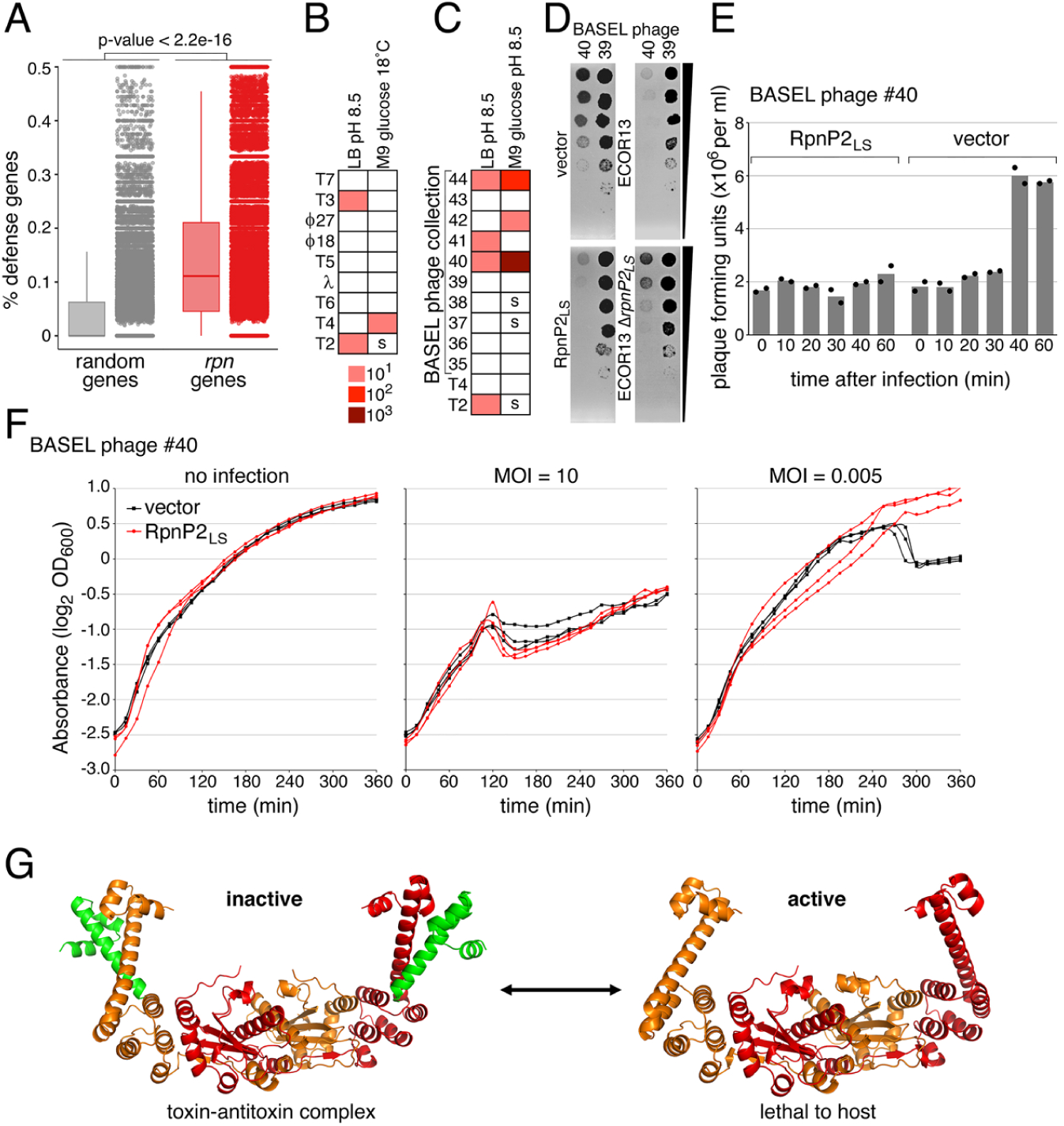
Rpn proteins block phage infection. (*A*) Enrichment of phage defense genes in the vicinity of *rpn* genes compared to random genes. Box plots (left) and dot plots (right) display the percentage and distribution of defense-associated genes in proximity to these genes, with P-values indicating the significance levels determined by the Mann-Whitney U Test. (*B*) Efficiency of plaquing (EOP) for the indicated phages infecting cells producing RpnP2_LS_ grown at 30°C on LB medium buffered to pH 8.5 and 18°C on M9 glucose medium pH 7.1. (*C*) EOP for the indicated phages infecting cells producing RpnP2_LS_ grown in LB and M9 glucose, pH 8.5. For (*B*) and (*C*), “s” denotes smaller plaques were observed. (*D*) Images of BASEL phage #40 and #39 plaques for MG1655 cells carrying vector control or producing RpnP2_LS_ (left) or ECOR13 or ECOR13 Δ*rpnP2_LS_* cells (right), from *SI Appendix,* Fig. S8. The back wedge denotes the 10-fold dilution series. (*E*) Growth of vector control or RpnP2_LS_ expressing cells infected with BASEL phage #40 (at indicated MOIs) in LB pH 8.5. Three biological replicates are presented. The partial regrowth of the strains infected at a MOI of 10 could be due to appearance of suppressors. (*F*) Plaque-forming units (PFU) per milliliter of BASEL phage #40 (MOI = 0.005) used to infect vector control or RpnP2_LS_ expressing cells at 0, 10, 20, 30, 40, and 60 min post-infection. Individual data points and average of two biological replicates are shown. For (*A*)-(*E*), the MG1655 strain background was WT for the *rpn* genes. **f**, Model for functions of Rpn_L_ and Rpn_S_ proteins. RpnA_L_ dimer (orange and red, right side) and RpnA_LS_ tetramer (left side, toxin-antitoxin complex) structures were predicted with AlphaFold-Multimer.

To test whether Rpn proteins constitute an anti-phage defense, we examined the ability of various phages to form plaques on *E. coli* MG1655 strains expressing plasmid-derived RpnP2_LS_ from its native promoter on a low copy plasmid (Fig. 5*B* and *C* and *SI Appendix*, Fig. S8). Considering the high levels of the RpnP2_S_ protein for cells grown in LB pH 7 (*SI Appendix*, Fig. S2*F*) could inhibit RpnP2_L_, we wanted to assay cells grown under conditions where the RpnP2_S_-RpnP2_L_ interaction might change. Given that RpnA_L*_ has higher activity at high pH and RpnA_LS_ complex tends to dissociate at high pH, we first grew cells in LB pH 8.5. Under these conditions, cells expressing RpnP2_LS_ showed 10-fold reduced plaquing by phage T2. We observed smaller T2 plaques and 10-fold reduced plaquing by T4 for RpnP2_LS_ expressing cells grown in minimal glucose medium pH 7.1 at low temperature (Fig. 5*B* and *SI Appendix*, Fig. S8*B*). We also examined the ability of other T-even phages in the BASEL phage collection (17) to plaque on this strain grown in both LB and minimal glucose media pH 8.5 (Fig. 5*C* and *SI Appendix*, Fig. S8*C* and *D*). RpnP2_LS_ protected against many of the T-even phages though resistance against BASEL phage #40 was the strongest (Fig. 5*D* and *SI Appendix*, Fig. S8*D*). This protection was mostly eliminated by active site mutations. Additionally, we deleted *rpnP*2 in the ECOR13 strain. This deletion strain showed reduced protection against BASEL phage #40 compared to the ECOR13 wt strain (Fig. 5*D SI Appendix*, Fig. S8*E*).

To further test for a phage defense function, we challenged RpnP2_LS_ expressing cells with BASEL #40 in liquid media at multiplicity of infections (MOIs) of either 10 or 0.005. Although both the vector control and RpnP2_LS_ cells were equally affected at high MOI, RpnP2_LS_ protects the cells at low MOI (Fig. 5*F*). A one-step growth curve also showed RpnP2_LS_ reduced the BASEL #40 burst size following a single round of infection (Fig. 5*E*). These data suggest Rpn_LS_ toxin-antitoxin systems can provide protection against specific phage.

## Discussion

Our data revealed that Rpn_S_ proteins are expressed together with Rpn_L_ proteins in *E. coli* as well as in *C. difficile* and serve as antitoxins for the toxic Rpn_L_ proteins (Fig. 5*G*). To our knowledge, this is the first time that a protein encoded within a gene has been characterized to be an antitoxin. It is possible that other toxin-antitoxin systems are comprised of additional components translated from iTIS as, for example, a clear ribosome profiling signal can be detected for *prlF* of the *yhaV-prlF* toxin-antitoxin system of *E. coli* (3, 4). Based on ribosome profiling data, genes-within-genes likely are far more prevalent than previously appreciated and not restricted to toxin-antitoxin genes.

Although the physiological roles of many toxin-antitoxin systems remain under debate, increasing numbers have been shown to constitute anti-phage defenses (12, 18–24). Our bioinformatics data showing *rpn* gene enrichment in defense islands as well as the variation in the four amino acid repeats in Rpn_S_ across different bacteria species were the initial hints that Rpn proteins also might protect against phage infection. Indeed, we demonstrated that a representative system, plasmid-encoded RpnP2_LS_, reduced plaque formation and infection by specific phages (Fig. 5*B*-*F*).

The intriguing variation in four amino acid repeats in Rpn_L_ and Rpn_S_ proteins is observed across a wide range of bacterial species. Changes in four amino acid repeats have been observed in the context of the EcoR124I and EcoR124II endonucleases (25). EcoR124I, with two repeats of the sequence “TAEL”, is specific for a sequence with a 6 bp gap (GAA**N_6_**RTCG), while EcoR124II, with three TAEL repeats, is specific for a sequence with a 7 bp gap (GAA**N_7_**RTCG). Our structural studies showed that RpnA_S_ is comprised of a three α-helix bundle. Additionally, HHPred (26) predicts relatedness to helix-turn-helix (HTH) motifs. In this context, it is noteworthy that many antitoxins have HTH motifs and adopt a strongly helical structure. In some examples, the toxin binding site overlaps these motifs (27, 28). The affinity between Rpn_S_ and Rpn_L_ is clearly tuned by the number of four amino acid repeats located in the long α-helix (Fig. 4*D*).

The overlapping *rpn_L_-rpn_S_* gene arrangement means there is obligatory co-evolution of the two proteins. The mechanism of the expansion and possibly contraction of these dodecamer repeats warrants further study. Given the extensive variation within a species, we suspect the changes ultimately are driven by a phage factor. It is possible that the Rpn_L_ proteins themselves are involved in increasing the number of four amino acid repeats as well as in initiating the recombination events that allow the generation of the small protein homologs encoded downstream of some homologs, and in generating the duplicated *rpn* genes.

How Rpn_L_ activity is released, a key unresolved issue for other toxin-antitoxin systems as well, is an important direction for future work. The dissociation of RpnA_L_ and RpnA_S_ and increased activity of RpnA_L_ at high pH, led us to wonder if Rpn proteins are activated at high pH or another specific physiological condition with similar effects. We found that RpnP2_LS_ provided more protection against different phages at alkaline pH. Since RpnA_S_ is stable at high pH, it is likely Rpn_S_ is released rather than degraded. In this context, we were intrigued to observe very different Rpn_L_:Rpn_S_ ratios depending on how these genes were expressed suggesting regulated expression is another factor in modulating Rpn_L_ activity. Other important questions are whether Rpn_L_ proteins have preferred substrates, whether the Rpn_L_ nucleases specifically recognize and cleave phage DNA, and what domains comprise the DNA-recognition determinants of the proteins. We expect that further mechanistic understanding of the actions of the Rpn_L_ and Rpn_S_ proteins will allow exploitation of these unique toxin-antitoxin systems.

## Materials and Methods

### Strains, Plasmids, and Oligonucleotides

Strains, plasmids, and oligonucleotides used in this study are listed in Dataset S2. Details about strain and plasmid construction are provided in *SI Appendix*.

### Bacterial Growth

Bacterial growth in Luria broth (LB) rich medium or M9 minimal medium was carried out at indicated pH, indicated temperature and indicated supplements as described in detail in *SI Appendix*.

### Immunoblot Analysis

Specific tagged proteins were detected by immunoblot analysis as described in detail in *SI Appendix*.

### β-Galactosidase Activity Assays

Induction of a *dinD-lacZ* reporter was assayed as described in detail in *SI Appendix*.

### Biochemical Characterization

RpnA protein purification and characterization by *in vitro* DNA cleavage assays, size exclusion chromatography (SEC) analysis and SEC-MALS analysis were carried out as described in detail in *SI Appendix*.

### Structure Determination and Prediction

The structure of RpnA_S_ was determined as described in detail in *SI Appendix* and as described in detail in *SI Appendix*. All Rpn proteins structures were predicted with AlphaFold 2.2.0 (13, 29) using NIH’s Biowulf cluster.

### Bioinformatic Analysis

Species distribution, phylogenetic analysis and gene context analysis were carried out as described in detail in *SI Appendix*.

### Bacteriophage Assays

Plaque assays and EOP measurements, growth curves following phage infection, and one-step growth curves to measure burst size were carried out as described in detail in *SI Appendix*.

### Data Availability

The data are available in the manuscript, or PDB 7TH0.

## ACKNOWLEDGMENTS

This work utilized the computational resources of the NIH HPC Biowulf cluster (http://hpc.nih.gov). We thank members of the Storz lab, A.S. Mankin and E. Raleigh for comments and A. Buskirk for generating browser images. Work by A.Z. was supported by an NICHD Early Career Award. This work was supported by the Intramural Programs of the *Eunice Kennedy Shriver* National Institute of Child Health and Human Development (A.Z., K.K., G.S.), National Library of Medicine (X.J.) and National Institute of Diabetes and Digestive and Kidney Diseases (A.H., F.D.), National Institutes of Health. M.T.L. is an Investigator of the Howard Hughes Medical Institute.

## Author contributions

A.Z., X.J., A.B.H., K.K., G.I.T.C, M.T.L., F.D. and G.S. designed experiments. A.Z., X.J., A.B.H., K.K. and F.D. performed experiments. A.Z., X.J., A.B.H., K.K., G.I.T.C, M.T.L., F.D. and G.S. analyzed data. A.Z., X.J., A.B.H., F.D. and G.S. wrote the paper.

