## Supplementary material for "Toxic anti-phage defense proteins inhibited by intragenic antitoxin proteins": SI Appendix

### Zhong et al.

**Strains, Plasmids, and Oligonucleotides.** The strains, plasmids, and oligonucleotides used in this study are listed in Dataset S2, respectively. Oligonucleotides were purchased from Integrated DNA Technologies. All the phages are from the lab stocks of M. Laub. BASEL phage collection (1) was originally shared by A. Harms. Rpn proteins were tagged on the chromosome following a published protocol (2). Specifically, the λ red recombination system in *E. coli* NM400 was used to replace the stop codon of each *rpn* gene, with the mutation linked to a SPA-kan cassette. After the replacement, the resulting constructs were moved into *E. coli* MG1655 by transduction using a lab stock of P1 phage. All constructs were verified by Sanger sequencing (Azenta Life Sciences). All cloning was performed using Q5® High-Fidelity 2X Master Mix (New England Biolabs, M0492L), NEBuilder® HiFi DNA Assembly Master Mix (New England Biolabs, E2621S) and NEB® Turbo Competent *E. coli* (New England Biolabs, C2984H). Site directed mutagenesis was performed with protocols from Agilent QuikChange II Site-directed mutagenesis kit with the exception that KOD Hot Start DNA polymerase (Sigma-Aldrich, 71086-3) and NEB® Turbo Competent *E. coli* were used. To eliminate the $P_{BAD}$ promotor and four transposition related genes, pMS34, pMS34-RpnA, pMS34-RpnB, and pMS34-RpnC (3) were modified to give pMS34*, pMS34*-$RpnA_{LS}$, pMS34*-$RpnB_{LS}$, and pMS34*-$RpnC_{LS}$. pEF21 was modified by the insertion of a ribosome binding site (RBS) to give pEF21* (4). pEF21* carrying $rpnA_S$, $rpnB_S$, $rpnC_S$, $rpnP2_{LS}$ and $rpnP2_{L*}$ were constructed with primers shown in Dataset S2. pBR322 was modified by removing the tetracycline promoter and resistance cassette to give pBR322*. $rpnA_{LS}$, $rpnB_{LS}$ and $rpnP2_{LS}$ genes together with the native promotors were cloned into pBR322*. SPA tag was inserted upstream of the stop codon of each *rpn* gene in pMS34*-RpnA$_{LS}$, pMS34*-RpnB$_{LS}$, pBR322*-RpnA$_{LS}$, pBR322*-RpnB$_{LS}$ and pBR322*-RpnP2$_{LS}$. For protein purification, the *rpn*$_{LS}$, *rpn*$_{L*}$, and *rpn*$_{S}$ genes were cloned into pTXB1 (New England Biolabs, N6707S). The genomic DNA for *C. difficile* strain 630 was purchased from American Type Culture Collection (ATCC, BAA-1382D-5). *E. coli* ECOR13 is from the lab stocks of M. Laub, but was originally obtained from the Thomas S. Whittam STEC Center at Michigan State University. To construct ECOR13 Δ*rpnP2*::*cm*, ECOR13 was first transformed with pKD46 (5). Primers with sequences flanking the *rpnP2* gene of the ECOR13 plasmid were used to amplify the $Cm^R$-resistance marker from pKD3. PCR products were electroporated into ECOR13 harboring pKD46. Colonies were screened for $Cm^R$ at 37˚C to cure the temperature sensitive pKD46 plasmid.

**Bacterial Growth.** Unless stated otherwise, bacterial cells were grown in Luria broth (LB) rich medium at 30˚C with shaking at 250 rpm. The *E. coli* strain ER2170 (New England Biolabs) was used in the growth experiments to parallel the DNA damage assays. *E. coli* ER2170 was transformed with pMS34*, pMS34*-RpnA$_{LS}$, pMS34*-RpnA$_{L*}$, pMS34*-RpnB$_{LS}$, pMS34*-RpnB$_{L*}$. A single colony was used to incubate a fresh LB medium containing 100 μg/mL Ampicillin (Amp). After overnight incubation at 30˚C, the culture was diluted into 10 mL of fresh LB with 100 μg/mL Amp to an $OD_{600}$ of 0.05, and the subculture was grown at 30˚C with shaking at 250 rpm. To ensure all the samples had the same $OD_{600}$, overnight cultures were first diluted to $OD_{600}$ ~ 0.6. After measuring the $OD_{600}$ for the diluted cultures, all the cultures were further diluted to $OD_{600}$ = 0.05. The same strategy was used for all the experiments involving diluting overnight cultures to $OD_{600}$ = 0.05. At 60 min, rhamnose (Thermo Fisher Scientific,

174081000) was added to a final concentration of 0.2%. After growing for another 4 h at 30˚C, the cultures were diluted to an $OD_{600}$ of 0.05 in LB with 100 µg/mL Amp and 0.2% rhamnose and 200 µL of the diluted cultures were dispensed in a 96-well plate (Falcon® 96-well Clear Flat Bottom TC-treated Culture Microplate). The plate was incubated in a CLARIOstar plate reader (BMG LABTECH) with temperature set at 30˚C. $OD_{600}$ was measured every 5 min with shaking at 300 rpm between each reading. For growth curve with RpnP2 in LB, *E. coli* MG1655 was transformed with pEF21*, pEF21*-$RpnP2_{LS}$ or pEF21*-$RpnP2_{L*}$. Glucose (0.4%) were added to the overnight cultures with 25 µg/mL chloramphenicol (Cm). Overnight cultures were washed with LB twice before diluting to $OD_{600}$ = 0.05 in LB (25 µg/mL Cm) with 0.4% glucose or 0.2% arabinose.

**Immunoblot Analysis.** To detect the SPA-tagged proteins, overnight cultures of the various strains were diluted into LB to a final $OD_{600}$ of 0.025. Cultures were sampled at around 2.5 and 5 h after dilution to represent the exponential growth phase ($OD_{600}$ of 0.5 to 0.7) and stationary growth phase ($OD_{600}$ of 2.5 to 3) respectively, and cells were collected by centrifugation at 4˚C. Cell numbers were normalized by $OD_{600}$. For 1.0 mL of culture, the cells were resuspended at 1 $OD_{600}$/80 µL in phosphate-buffered saline. The suspended cells were mixed with 2x Laemmli sample buffer (Bio-Rad, 1610737), heated at 95˚C for 10 min and centrifuged at 4˚C for 3 min. The resulting supernatant was analyzed by SDS-PAGE (Bio-Rad, 4561093, 4–20%). The gel was transferred to nitrocellulose membranes (Invitrogen) and probed with anti-FLAG(M2)-HRP (Sigma-Aldrich, A8592-1MG) according to the manufacturer's protocol. For the detection of SPA-tagged proteins upon rhamnose induction in the MG1655 background, 0.2% rhamnose was added to cells and samples were collected at ~2.5 h. For the detection of SPA-tagged proteins with rhamnose induction in ER2170 background, overnight cultures were diluted to $OD_{600}$ = 0.05. At 60 min, rhamnose was added to a final concentration of 0.2%. After growing for another 4 h at 30˚C, the cultures were diluted to an $OD_{600}$ of 0.05 in LB with 100 µg/mL amp and 0.2% rhamnose and samples were collected around 6 h.

*rpn* gene (protein id: WP_009902002.1) was amplified from *C. difficile* strain 630 genomic DNA and cloned into pTXB1 vector. Two different mutants to disrupt the predicted $Rpn_S$ RBS and iTIS, $Rpn_{L*\text{-}M1}$ and $Rpn_{L*\text{-}M2}$, were generated with NEBuilder® HiFi DNA Assembly Master Mix using oligonucleotides listed in Dataset S2. The resulting plasmids were transformed into *E. coli* T7 Express *lysY/I$^q$* (New England Biolabs, C3013I). Six colonies were inoculated into 5 mL of LB with 100 µg/mL Amp. The culture was incubated at 30˚C for 4.5 h and was further diluted 12-fold in 30 mL of LB with 100 µg/mL Amp. Upon reaching $OD_{600}$ ~ 0.6, the culture was cooled at 4˚C for 1 h. Isopropyl β-D-thiogalactoside (IPTG) was added to the culture to a final concentration of 0.4 mM. The culture then was incubated at 15˚C with shaking at 100 rpm for another 20 h. Cells were collected by centrifugation at 4,000 x g for 10 min at 4˚C. After discarding the supernatant, the cells were resuspended in 2 mL buffer containing 20 mM Tris-HCl, 500 mM NaCl, pH 8.0. The cells were lysed with sonication (Branson 450 Digital Sonifier). The resulting mixture was centrifuged at 11,000 x g for 10 min at 4˚C. The samples were diluted 20-fold before loading. The diluted samples were analyzed by SDS-PAGE (4–20%). TCEP (45 mM) instead of β-mercaptoethanol was added to the sample buffer as the reducing agent to minimize the cleavage of the intein tag during sample preparation. The gel was transferred to nitrocellulose membranes and probed with anti-CBD monoclonal antibody (New England Biolabs, E8034S) as the primary antibody (10 µL of antibody in 10 mL 5% of non-fat milk at 4˚C overnight) and goat anti-mouse IgG HRP-conjugated antibody (Pierce, 1858413) as the secondary antibody (3 μL of antibody in 15 mL of 5% non-fat milk at 4˚C overnight, according to the manufacturer's protocol).

**β-Galactosidase Activity Assays.** For some of the assays, *E. coli* ER2170 was transformed with one plasmid, pMS34*, pMS34*-$RpnA_{LS}$, or pMS34*-$RpnB_{LS}$. The $RpnA_{L*}$ and $RpnB_{L*}$ derivatives could not be used in these assays because of their toxicity. To obtain the two-plasmid strains, *E. coli* ER2170 harboring pMS34*-$RpnA_{LS}$ was transformed with pEF21*, pEF21*-$RpnA_{S}$, pEF21*- $RpnA_{S\text{-}STOP}$ or pEF21*-$RpnB_{S}$, ER2170 harboring pMS34*-$RpnB_{LS}$ was transformed with pEF21*, pEF21*-$RpnB_{S}$, pEF21*-$RpnB_{S\text{-}STOP}$ or pEF21*- $RpnA_{S}$, or ER2170 harboring pMS34*-$RpnC_{LS}$ was transformed with pEF21*, pEF21*-$RpnC_{S}$ or pEF21*-$RpnC_{S+REPEAT}$. For the experiments with $RpnC_{LS+REPEAT}$, ER2170 was simultaneously transformed with pMS34*-$RpnC_{LS+REPEAT}$ and pEF21*, pEF21*-$RpnC_{S}$ or pEF21*-$RpnC_{S+REPEAT}$.

For single-plasmid experiments with RpnA and RpnB, a single colony was used to incubate a fresh RB medium (10 g/L tryptone, 5 g/L yeast extract, 5 g/L NaCl, pH 7.2) containing 100 μg/mL Amp. After overnight incubation at 30˚C, the culture was diluted into 10 mL of fresh RB with 100 μg/mL Amp to an $OD_{600}$ of 0.05, and the subculture was grown at 30˚C. At 1.0 h, rhamnose was added to a final concentration of 0.2% (to induce $Rpn_{L}$ expression). For each sample, 100 μL of culture was taken at 7.5 h for the β-galactosidase activity assay. $OD_{600}$ was recorded at the same time.

For two-plasmid experiments with RpnA and RpnB, a single colony was used to incubate a fresh RB medium containing 100 μg/mL Amp and 25 μg/mL Cm. After overnight incubation at 30˚C, the culture was diluted into 10 mL of fresh RB with 100 μg/mL Amp and 25 μg/mL Cm to an $OD_{600}$ of 0.05, and the subculture was grown at 30˚C. At 1.0 h, arabinose (Sigma-Aldrich, W325501-100G) was added to a final concentration of 0.2% (to induce $Rpn_S$ expression). At 2.0 h, rhamnose was added to a final concentration of 0.2% (to induce $Rpn_L$ expression). Aliquots (100 µL) were taken at 9.0 h for the RpnA samples and 7.5 h for the RpnB samples based on growth curves in (3). $OD_{600}$ was recorded at the same time.

For two-plasmid experiments with RpnC, after overnight incubation in LB at 30˚C, the culture was diluted into 10 mL of fresh RB with 100 µg/mL Amp and 25 µg/mL Cm to an $OD_{600}$ of 0.05, and the subculture was grown at 23˚C. At 2.0 h, arabinose was added to a final concentration of 0.2% (to induce $RpnC_S$ expression) and rhamnose was added to a final concentration of 0.2% (to induce $RpnC_L$ expression). Aliquots (100 µL) were taken at 16 h. $OD_{600}$ was recorded at the same time.

For each batch of assays, 30 mL of reaction buffer was freshly prepared by mixing 30 mL of Z buffer (60 mM $Na_2HPO_4$, 40 mM $NaH_2PO_4$, 10 mM KCl and 1 mM $MgSO_4$) with 45 µL of 0.1% SDS and 81 µL β-mercaptoethanol. To a 1.5 mL Eppendorf tube containing 100 µL of previously collected culture, 700 µL of reaction buffer and 30 µL of chloroform were added. After vortexing for 30 s, all the samples were incubated at 28˚C for 15 min. Subsequently, 100 µL of *o*-nitrophenyl β-D-galactopyranoside (Sigma-Aldrich, N1127-5G, ONPG, 8 mg/mL) was added to each tube. The reaction tubes were incubated at 28˚C for another 7-60 min. All the reactions were quenched by adding 500 µL of 1.0 M $Na_2CO_3$. After centrifugation at 14,000 rpm for 3 min, $OD_{420}$ was recorded for the supernatant. Miller units were calculated as reported (6). Three or four independent biological replicates were performed for all the assays.

**Rpn Protein Characterization**

**Protein Overexpression and Purification.** Protein purifications were achieved with a hand-packed gravity column or a prepacked column on a AKTA Purifier 10 FPLC system (Cytiva). pTXB1 derivatives carrying *rpnA*$_{LS}$ and *rpnA*$_{S}$ were transformed into T7 Express competent *E. coli* (New England Biolabs, C2566H). pTXB1 derivatives carrying *rpnA*$_{L*}$ was transformed into *E. coli* T7 Express *lysY/I*$^{q}$ (with *rpnA* gene deleted). Six colonies were inoculated into 10 mL of LB with 100 µg/mL Amp. The overnight culture was diluted 10-fold to a fresh LB culture with 100 µg/mL Amp. After growing for 1 h, the culture was further diluted 25-fold in 1.0 L of LB with 100 µg/mL Amp. Upon reaching $OD_{600}$ ~ 0.6, the culture was cooled at 4˚C for 2 h. IPTG was added to the culture to a final concentration of 0.4 mM. The culture then was incubated at 15˚C with shaking at 100 rpm for another 20 h. Cells were collected by centrifugation at 5,000 x g for 15 min at 4˚C. After discarding the supernatant, the cells were resuspended in buffer A (20 mM Tris-HCl, 500 mM NaCl, 0.5 mM TCEP, pH 8.0) with 50 mL buffer per 10 g cells. The cells were lysed with sonication. The resulting mixture was centrifuged at 15,000 x g for 30 min at 4˚C. A small fraction of the supernatant was subject to immunoblot analysis. The samples corresponding to RpnA$_{LS}$ and RpnA$_{L*}$ were diluted 10-fold before loading. RpnA$_{S}$ sample was diluted 40-fold. The diluted samples were analyzed by SDS-PAGE (4–20%). The gel was transferred to nitrocellulose membranes and blotted against anti-CBD monoclonal antibody as the primary antibody (10 µL of antibody in 10 mL 5% non-fat milk at 4˚C overnight) and goat anti-mouse IgG HRP-conjugated antibody (3 µL of antibody in 15 mL 5% non-fat milk at 4˚C overnight) as the secondary antibody according to the manufacturer's protocol. The rest of the supernatant was applied to a gravity column with 10 mL of pre-washed chitin resin (New England Biolabs, S6651L). After washing with 500 mL of buffer A, 60 mL of cleavage buffer (buffer A with 50 mM β-mercaptoethanol) was applied to the column rapidly. The column was maintained at 4˚C for 40 h to maximize the cleavage of intein tag. The desired protein was eluted with buffer A and concentrated with Amicon® Ultra-15 centrifugal filter (Sigma-Aldrich).

RpnA$_{LS}$ and RpnA$_{L*}$ were further purified by anion-exchange chromatography. RpnA$_{LS}$ and RpnA$_{L*}$ were buffer-exchanged into buffer B (20 mM Tris-HCl, 100 mM NaCl, 0.5 mM TCEP, pH 8.0) and loaded onto a HiTrap Q HP 5-mL column (Cytiva). The protein was eluted with a 150 mL-linear gradient (from 100 mM to 600 mM NaCl). The desired fractions were pooled and concentrated. RpnA$_S$ was further purified by size-exclusion column. Concentrated RpnA$_S$ was directly load onto a HiLoad 16/60 Superdex 75 prep grade column (Cytiva). The protein was eluted with an isocratic condition (buffer B). The desired fractions were pooled and concentrated. The purity of RpnA$_{LS}$, RpnA$_{L*}$ and RpnA$_S$ were determined by SDS-PAGE analysis. Purified RpnA$_{LS}$, RpnA$_{L*}$ and RpnA$_S$ were stored at -80˚C. We note that attempts to overexpress and purify RpnB proteins failed due to higher toxicity.

To generate selenomethionine-substituted RpnA$_S$ (SeMet RpnA$_S$), pTXB1 derivatives carrying *rpnA$_S$* was freshly transformed into B834(DE3) Competent Cells (Novagen). Six colonies were inoculated into 100 mL of SelenoMethionine Medium Base (Molecular Dimensions, MD12-501) with SelenoMethionine Medium Nutrient Mix (Molecular Dimensions, MD12-502), 40 µg/mL methionine, and 100 µg/mL Amp. The overnight culture was pelleted, washed three times with 100 mL of water, and resuspended in 4.8 mL of water. Then, 0.8 mL of the resuspended cells was used to inoculate 1 L of SelenoMethionine Medium Base with SelenoMethionine Medium Nutrient Mix, 40 µg/mL selenomethionine, and 100 µg/mL Amp. After growing for about 7 h, $OD_{600}$ reached ~ 0.5. The culture was cooled at 4˚C for 2 h. IPTG was added to the culture to a final concentration of 0.4 mM. The culture was then incubated at

15°C with shaking at 100 rpm for another 20 h. SeMet $RpnA_S$ was purified in the same way as $RpnA_S$.

**In Vitro DNA Cleavage Assays.** To normalize protein concentration, $RpnA_{LS}$ and $RpnA_{L*}$ concentrations were determined using the calculated extinction coefficient of RpnA ($\varepsilon_{280}$ = 27390 $M^{-1}\cdot cm^{-1}$, calculated with ProtParam tool by Expasy) (7). As $RpnA_S$ does not have any tryptophan or tyrosine resides and only one phenylalanine, $RpnA_S$ concentration was estimated based on the extinction coefficient of phenylalanine ($\varepsilon_{257.5}$ = 195 $M^{-1}\cdot cm^{-1}$). In a total volume of 20 µL, purified $RpnA_{L*}$, $RpnA_{LS}$ or $RpnA_S$ (9.6 µM) was incubated with 50 ng/µL of pUC19 (New England Biolabs, N3041S) or M13MP18 ssDNA (New England Biolabs, N4040S) in 50 mM Tris-HCl, pH 8.0 at 37 °C for 6 h. To determine the effect of $RpnA_S$ on the DNA endonuclease activity of $RpnA_{L*}$, 9.6 µM (1X) or 19.2 µM (2X) $RpnA_S$ was added to $RpnA_{L*}$ (9.6 µM) assay with pUC19 dsDNA or M13MP18 ssDNA. Purified $RpnA_{L*}$ or $RpnA_{LS}$ (9.6 µM) was incubated with pUC19 (50 ng/µL) in 50 mM HEPES (pH 7.0), 50 mM Tris-HCl (pH 8.0) or 50 mM CHES (pH 9.0) at 37 °C in a total volume of 50 µL for 6 h. All the assays also contain 50 mM NaCl, 10 mM $MgCl_2$, 15 mM $CaCl_2$ and 1 mM DTT. All the assays were quenched by addition of EDTA to the final concentration of 83 mM and subsequently analyzed by 1.0 % agarose gel running at 100 V.

**Size Exclusion Chromatography (SEC) Analysis.** $RpnA_{LS}$ and $RpnA_{L*}$ concentrations were determined by Bradford assay. The concentrations obtained were very close to the ones estimated by extinction coefficient. Bradford assay could not be used to estimate the concentration for $RpnA_S$ because there are not enough positive charges on $RpnA_S$. Thus, $RpnA_S$

concentration was estimated based on the extinction coefficient of phenylalanine. All experiments were performed with a Superdex 200 Increase 10/300 GL column (Cytiva) or a Superdex 75 Increase 10/300 GL column (Cytiva) in buffer B. For $RpnA_{LS}$, 200 µL of 0.5 mg/mL protein was injected. For $RpnA_S$, 160 µL of 10.2 mg/mL protein was injected. To determine if $RpnA_{L*}$ and $RpnA_S$ interact with each other, 150 µL of 3.0 mg/mL $RpnA_{L*}$ was incubated with 150 µL of 10.2 mg/mL $RpnA_S$ at room temperature for 30 min before injection (Fig. 4*A*). For *SI Appendix,* Fig. S5*A*, 150 µL of 3.0 mg/mL $RpnA_{L*}$ was incubated with 150 µL of 10.4 mg/mL $RpnA_S$ at room temperature for 30 min before injection. Fractions were analyzed by tris-glycine SDS-PAGE. As a control, 150 µL of 3.0 mg/mL $RpnA_{L*}$ was incubated with 150 µL buffer B for 30 min at room temperature before injection. For *SI Appendix,* Fig. S5*B*, $RpnA_{LS}$ (0.7 mg/mL) was incubated in 100 mM HEPES (pH 7.0), 100 mM Tris-HCl (pH 8.0), 100 mM CHES (pH 9.0) or 100 mM CAPS (pH 10.0) together with 0.1 M NaCl and 0.5 mM TCEP in a total volume of 250 µL at room temperature overnight.

**SEC-MALS Analysis.** Experiments were carried out in an Agilent Series 1100 System coupled with a Optilab T-rEX refractive index (RI) detector (Wyatt Technology) and a Heleos-II in-line multi-angle light scattering (LS) detector (Wyatt Technology). For $RpnA_S$, Superdex 75 Increase 10/300 GL column was used. For $RpnA_{LS}$, Superdex 200 Increase 10/300 GL column was used instead. All the experiments were carried out at room temperature. The system was washed with buffer B for approximately one hour until a stable baseline for the RI detector was reached. SEC columns were washed for another hour without connecting to the RI and LS detectors. The system was equilibrated again after connecting the SEC column to the detectors. All samples were analyzed at a flow rate of 0.5 ml/min with buffer B. Before running $RpnA_S$ and $RpnA_{LS}$, 50 μL of 2 mg/mL BSA (Thermo Fisher Scientific, 23209) was injected as a standard. For $RpnA_S$, 50 μL of 14.5 mg/mL sample was injected. For $RpnA_{LS}$, 50 μL of 3.1 mg/mL sample was injected. All data were analyzed with ASTRA software (V7.1, Wyatt Technology).

**Structure Determination and Prediction**

**Structure Determination.** Protein crystals were obtained through hanging drop method. A drop of 2 μL of SeMet $RpnA_S$ (8.6 mg/mL) in buffer B was added to 2 μL crystallization reagent (0.5 M LiCl, 0.1 M Tris-HCl, pH 8.3, 8.6 or 9.0, 30 % PEG 6,000). Then, 0.2 or 0.3 μL 0.1 M Iron(III) chloride hexahydrate was added before sealing the cover slip. Crystals reached maximum size after 3 days at 20˚C. Crystals were harvested by freezing with liquid nitrogen directly.

The structure was solved with experimental phasing using a Se-Met substituted protein. A highly redundant (7.4-fold, Friedel's law false) data set was collected on a single crystal in two 200º rotation sweeps at different goniometer arc settings. There was no appreciable radiation decay. Data were collected using 1.5418 Å radiation from a Rigaku FR-X X-ray source and an Eiger2 4M pixel array detector at 95K. Data were integrated and scaled using XDS and XSCALE (8). Single wavelength Anomalous Scattering (SAS or SAD) phasing and density modification were carried out using Phenix's (9) AutoSol procedure that resulted in a clearly interpretable electron density map. Phenix's autobuilt model was completed manually using O (10), and was refined using energy minimization in Phenix. The final model contains residues between M244 and A289; the three C-terminal residues are disordered. 100% of the residues are in the most favored region of the Ramachandran plot. Interfaces were analyzed using 'Protein interfaces, surfaces and assemblies' service PISA (11) at the European Bioinformatics Institute (http://www.ebi.ac.uk/pdbe/prot_int/pistart.html). Information about the structure is given in *SI Appendix*, Table S1.

**Structure Prediction.** All Rpn proteins structures were predicted with AlphaFold 2.2.0 (12, 13) using NIH's Biowulf cluster. The predictions with the highest confidence were shown. pLDDT scores were shown in different colors to represent the model confidence.

**Bioinformatic Analysis**

**Analysis of Apecies Distribution.** To investigate the species distribution of the *rpn* orthologs, we searched the representative genomes in Genome Taxonomy Database (GTDB release202) (14-16). Gene prediction and annotation were performed using Prokka (17) (version 1.14.6). Clusters of Orthologous Group (COG) annotation for genes was determined using eggNOG-mapper (version 2.1.6) with eggNOG orthology data (version 5.0.2) (18). Genes annotated with COG ID "COG5464" were identified as *rpn* orthologs. The identified Rpn genes, along with their sequences, taxonomic distribution, domain annotation and Neighboring genes, are available in Dataset S1 and the Github link(https://github.com/nlm-irp-jianglab/rpn_data). The taxonomic distribution of *rpn* orthologs was plotted with iTOL (15). The results were plotted at the taxonomic level of order and above. If all children of a clade have the same *rpn* presence/absence status, only the upper level is displayed.

**Phylogenetic Analysis.** To construct the phylogenetic tree of ECOR genomes (19), we utilized the Roary (20) software (version 3.13.0) with parameters set at "-i 90 -cd 90 -s -e –mafft" to extract the core gene alignments. These alignments were then used to build the tree using

FastTree (version 2.1.10) with parameters set at "-nt -gtr". For Rpn protein sequences, alignments were performed with Clustal-Omega (version 1.2.4) with the default parameters. The phylogenetic tree was built from the alignments using FastTree (version 2.1.10 ) with default parameters (21). iTOL (15) was used to visualize the phylogenetic tree. The ECOR Rpn alignments and their phylogenetic trees were provided in a Github repository (https://github.com/nlm-irp-jianglab/rpn_data).

**Gene Context Analysis.** We conducted genomic context analysis using a method similar to the one described previously (22). We randomly selected an equal number of random and *rpn* genes and identified the flanking genes for both sets of genes. The local genomic context of a gene was defined as all coding sequences within a region of ±10 kb of the gene or to the end of a contig if less than 10 kb. We used HMMER3 hmmscan (version 3.3.2) with E-value cutoffs of $10^{-5}$ to search for defense-related domains in all coding sequences within this local context (23). Specifically, we searched for defense-related Pfam and COG domains that were previously identified in studies by Makarova et al. (24), Doron et al. (25), and Gao et al. (26).

**Bacteriophage Assays**

**Plaque Assays and EOP Measurements.** Plaque assays and EOP measurements were carried out based on published protocols (22, 27-29). *E. coli* MG1655 was transformed with pBR322*, pBR322*-RpnP2$_{LS}$ and pBR322*-RpnP2$_{LS+M}$. For plaque assays with these strains, 100 µL of a bacterial strain of interest was mixed with 4.6 mL LB pH 8.5 (50 mM Tris, 2 mM $MgSO_4$ and 0.1 mM $CaCl_2$) or M9-glucose pH 8.5 (50 mM Tris, 1.1 g/L $Na_2HPO_4 \cdot 7H_2O$, 0.26 g/L $KH_2PO_4$, 0.25 g/L NaCl, 0.5 g/L $NH_4Cl$, 0.1% casamino acids, 0.4% glucose, 2 mM $MgSO_4$ and 0.1 mM

$CaCl_2$) + 0.5% agar and spread on plates with 40 mL of LB pH 8.5 or M9 glucose pH 8.5 plus 1.2% agar and 100 µg/mL carbenicillin. For plaque assays with the ECOR13 strains, carbenicillin was omitted. Tenfold serial diluted phages (2 µL) were spotted on the bacterial lawn. Images were taken after overnight incubation at 30˚C. For plaque assays at 18˚C, 100 µL of a bacterial strain of interest was mixed with 4.6 mL M9-glucose pH 7.1 (6.4 g/L $Na_2HPO_4 \cdot 7H_2O$, 1.5 g/L $KH_2PO_4$, 0.25 g/L NaCl, 0.5 g/L $NH_4Cl$, 0.1% casamino acids, 0.4% glucose, 2 mM $MgSO_4$ and 0.1 mM $CaCl_2$) + 0.5% agar and spread on plates with M9 glucose pH 7.1, 1.2% agar, and 100 µg/mL carbenicillin. All phage lysate stocks were prepared with *E. coli* MG1655 according to published protocols (30). Phage titers were determined by mixing serial dilutions with top agar (31).

**Grow Curve Following Phage Infection in Liquid Media.** *E. coli* MG1655 harboring pBR322* or pBR322*-RpnP2$_{LS}$ were grown in LB pH 8.5. Overnight cultures were diluted to $OD_{600}$ around 0.2 and 180 µL of the diluted cultures were dispensed to a 96-well plate. Then, 20 µL of phage in different concentrations were added to reach different MOI. The plate was incubated in a CLARIOstar plate reader at 30˚C with shaking at 300 rpm. $OD_{600}$ was measured every 15 min.

**One-Step Growth Curves to Measure Burst Size.** One-step growth curves were performed following published procedures (29, 32). *E. coli* MG1655 harboring pBR322* or pBR322*-RpnP2$_{LS}$ were grown in LB pH 8.5. Overnight cultures were diluted to $OD_{600}$ around 0.1. After reaching $OD_{600}$ around 0.4, 450 µL cells were infected with phage at an MOI of 0.005. Phages were allowed to adsorb to the cells for 10 min before diluting 10,000-fold. Cells were continually shaking at 30˚C. At each time point, 100 or 200 μL of the infected cells were taken and mixed with MG1655 strain. The mixture was mixed with 3 mL LB 0.5% agar and spread on 25 mL LB 1.2% agar plate. Plaques were counted after incubated at 37˚C overnight. Data reported were the mean of two biological repeats with individual data points shown.

1. E. Maffei *et al.*, Systematic exploration of Escherichia coli phage-host interactions with the BASEL phage collection. *PloS Biol.* **19**, e3001424 (2021).
2. D. Yu *et al.*, An efficient recombination system for chromosome engineering in *Escherichia coli*. *Proc. Natl. Acad. Sci. USA* **97**, 5978-5983 (2000).
3. A. W. Kingston, C. Ponkratz, E. A. Raleigh, Rpn (YhgA-Like) proteins of *Escherichia coli* K-12 and their contribution to RecA-independent horizontal transfer. *J. Bacteriol.* **199**, pii: e00787-00716 (2017).
4. E. M. Fozo *et al.*, Repression of small toxic protein synthesis by the Sib and OhsC small RNAs. *Mol. Microbiol.* **70**, 1076-1093 (2008).
5. K. A. Datsenko, B. L. Wanner, One-step inactivation of chromosomal genes in *Escherichia coli* K-12 using PCR products. *Proc. Natl. Acad. Sci. USA* **97**, 6640-6645 (2000).
6. J. H. Miller, *A Short Course in Bacterial Genetics: A Laboratory Manual and Handbook for Escherichia coli and Related Bacteria.* (Cold Spring Harbor Laboratory Press, Plainview, NY, 1992).
7. Y.-b. Lim, E. C. Lee, M. Lee, Cell-penetrating-peptide-coated nanoribbons for intracellular nanocarriers. *Angew. Chem. Int. Ed. Engl.* **46**, 3475-3478 (2007).
8. K. W., XDS. *Acta Crystallogr D Biol Crystallogr.* **66**, 125-132 (2010).
9. T. C. Terwilliger *et al.*, Decision-making in structure solution using Bayesian estimates of map quality: the PHENIX AutoSol wizard. *Acta Crystallogr D Biol Crystallogr.* **65**, 582-601 (2009).
10. T. C. Terwilliger, O. V. Sobolev, P. V. Afonine, P. D. Adams, Automated map sharpening by maximization of detail and connectivity. *Acta Crystallogr D Struct Biol.* **74**, 545-559 (2018).
11. E. Krissinel, K. Henrick, Inference of macromolecular assemblies from crystalline state. *J. Mol. Biol.* **372**, 774-797 (2007).
12. J. Jumper *et al.*, Highly accurate protein structure prediction with AlphaFold. *Nature* **596**, 583-589 (2021).
13. R. Evans *et al.*, Protein complex prediction with AlphaFold-Multimer. *bioRxiv*, 10.1101/2021.1110.1104.463034 (2021).
14. D. H. Parks *et al.*, Recovery of nearly 8,000 metagenome-assembled genomes substantially expands the tree of life. *Nat. Microbiol.* **2**, 1533–1542 (2017).
15. I. Letunic, P. Bork, Interactive Tree Of Life (iTOL) v4: recent updates and new developments. *Nucleic Acid Res.* **47**, W256-W259. (2019).
16. D. H. Parks *et al.*, Author Correction: A complete domain-to-species taxonomy for Bacteria and Archaea. *Nat. Biotechnol.* **38**, 1098 (2020).

**Table S1.** Data collection and refinement statistics.

| | Crystal 1<br>$RpnA_S$ |
|---|---|
| **Data collection** | |
| Space group | C222 (1) |
| Cell dimensions | |
| *a*, *b*, *c* (Å) | 38.04 58.29<br>38.72 |
| α, β, γ (°) | 90.0, 90.0, 90.0 |
| Resolution (Å) | 30.0-1.9 (1.95-<br>1.90) |
| $R_{merge}$ | 0.029 (0.413) |
| $I / \sigma I$ | 43.46 (6.87) |
| Completeness (%) | 99.8 (100.0) |
| Redundancy | 13.38 (11.69) |
| **Refinement** | |
| Resolution (Å) | 19 – 1.9 |
| No. reflections | 3582 |
| $R_{work} / R_{free}$ | 0.226/0.247 |
| No. atoms | |
| Protein | 353 |
| Ligand/ion | 0 |
| Water | 8 |
| *B*-factors | |
| Protein | 55.05 |
| Ligand/ion | |
| Water | 56.12 |
| R.m.s deviations | |
| Bond lengths (Å) | 0.007 |
| Bond angles (°) | 0.793 |

*Values in parentheses are for highest-resolution shell.

PDB ID code 7TH0

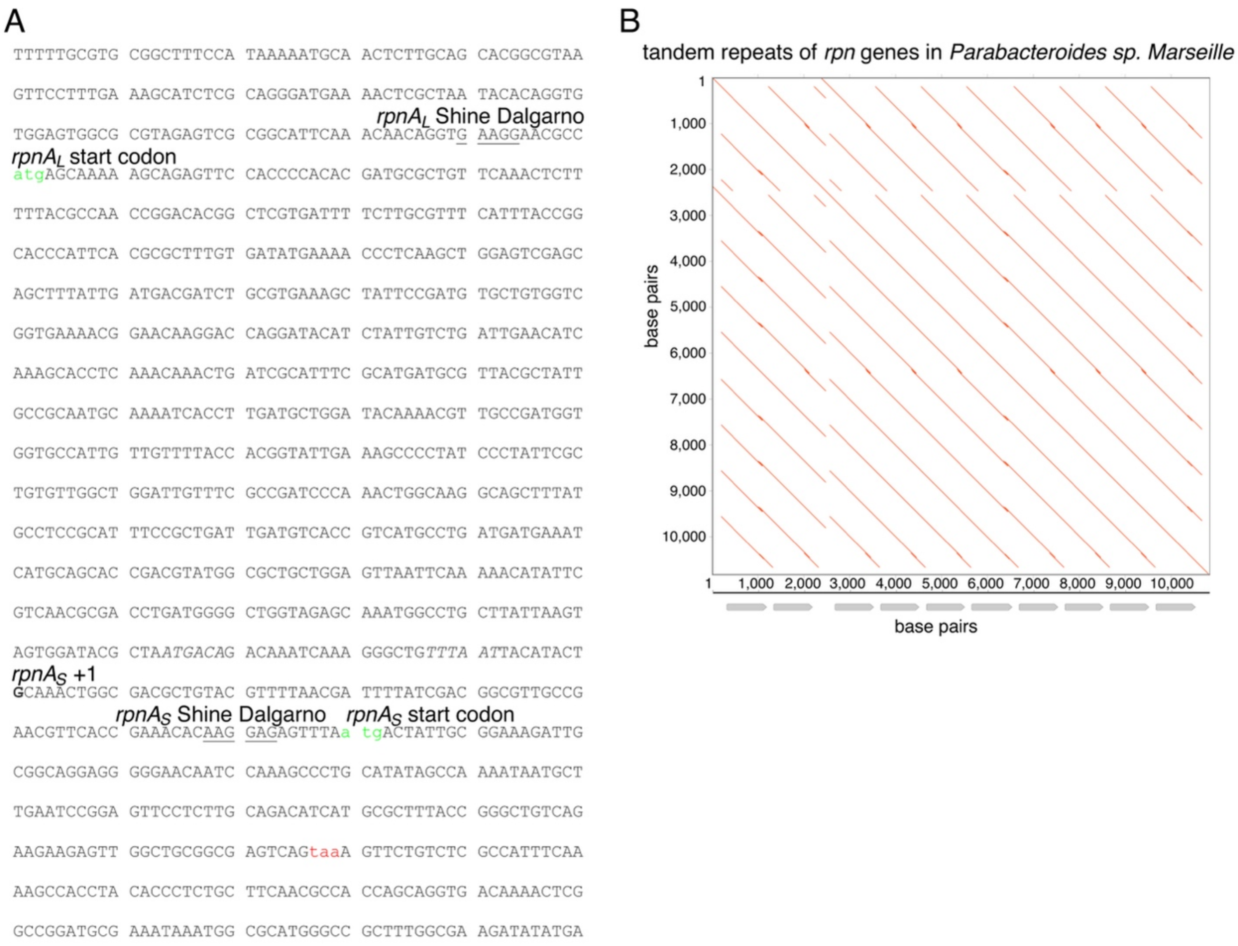

**Fig. S1.** Genomic sequence and predicted domain structure of $RpnA_L$ protein and tandem repeats of *rpn* genes. (*A*) The sequence of the *E. coli rpnA* gene with Shine Dalgarno sequences (underlined) and start codons (green font) indicated for both $RpnA_L$ and $RpnA_S$. (*B*) The *rpn* genes (grey arrows) in *Parabacteroides sp. Marseille* are in a tandem repeat array. The dot plot was generated using self-aligned DNA sequences of the *rpn* gene array locus. The tandem repeats were represented as a series of parallel diagonals whose frequency and spacing reflect their abundance and unit lengths.

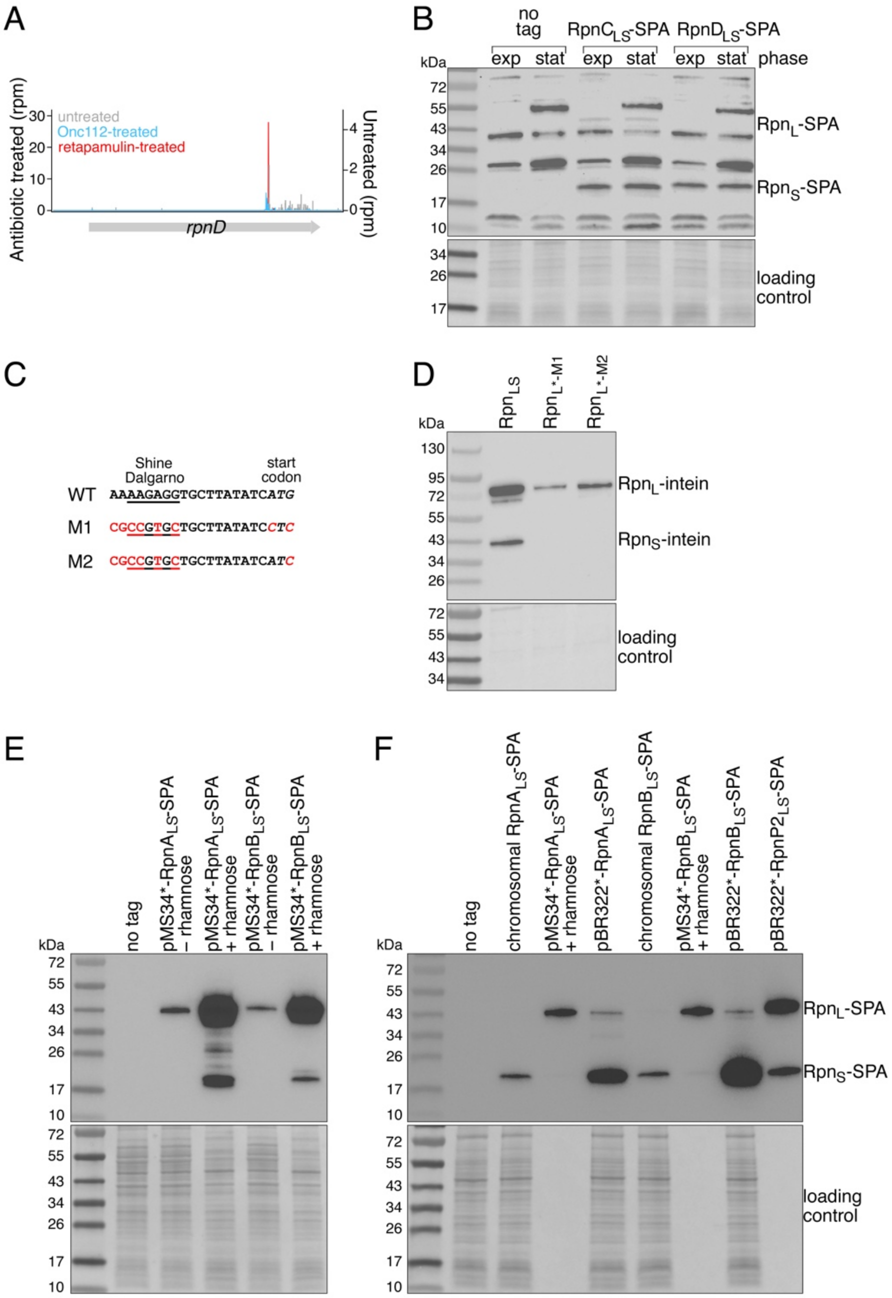

**Fig. S2.** Small proteins ($\mathrm{Rpn_S}$) encoded within the *C*-terminal domain of $\mathrm{Rpn_L}$ are synthesized in diverse organisms. (*A*) Browser images of ribosome profiling data for *rpnD*, which is a pseudogene in *E. coli* K-12. (*B*) Immunoblot analysis of $\mathrm{RpnC_S}$ and $\mathrm{RpnD_S}$ levels. Assay was carried out as in Fig. 1. A stop codon is present in the middle of *E. coli* MG1655 *rpnD*, a pseudogene. Thus, $RpnD_L$ was not detected. (*C*) Mutations introduced in the Shine-Dalgarno sequence and start codon for *C. difficile* $Rpn_S$. (*D*) Immunoblot analysis of *C. difficile* $Rpn_L$-intein and $Rpn_S$-intein expressed from $Rpn_{LS}$, $Rpn_{L*-M1}$ and $Rpn_{L*-M2}$. Amino acid substitutions necessitated by Shine-Dalgarno and start codon mutations reduce the levels of the $Rpn_L$ protein by affecting translation or stability. (*E*) Immunoblot analysis of the levels of pMS34*-$RpnA_{LS}$-SPA and pMS34*-$RpnB_{LS}$-SPA without and with rhamnose induction in a ER2170 strain background. (*F*) Immunoblot analysis of the levels of chromosomally SPA-tagged $RpnA_S$; pMS34*-$RpnA_{LS}$-SPA with rhamnose induction (sample was diluted 200-fold); pBR322*-$RpnA_{LS}$-SPA with native *rpnA* promotor; chromosomally SPA-tagged $RpnB_S$; pMS34*-$RpnB_{LS}$-SPA with rhamnose induction (sample was diluted 200-fold); pBR322*-$RpnB_{LS}$-SPA with native *rpnB* promotor; pBR322*-$RpnP2_{LS}$-SPA with native *rpnP2* promotor (sample was diluted 1,000-fold) in an MG1655 strain background. For (*B*), (*E*), and (*F*), the SPA tag was detected with monoclonal anti-FLAG M2-peroxidase (HRP) antibody, while for (*D*), the intein tag was detected with anti-CBD monoclonal antibody. For (*B*), (D), (*E*), and (*F*), Ponceau S staining of membranes served as loading controls.

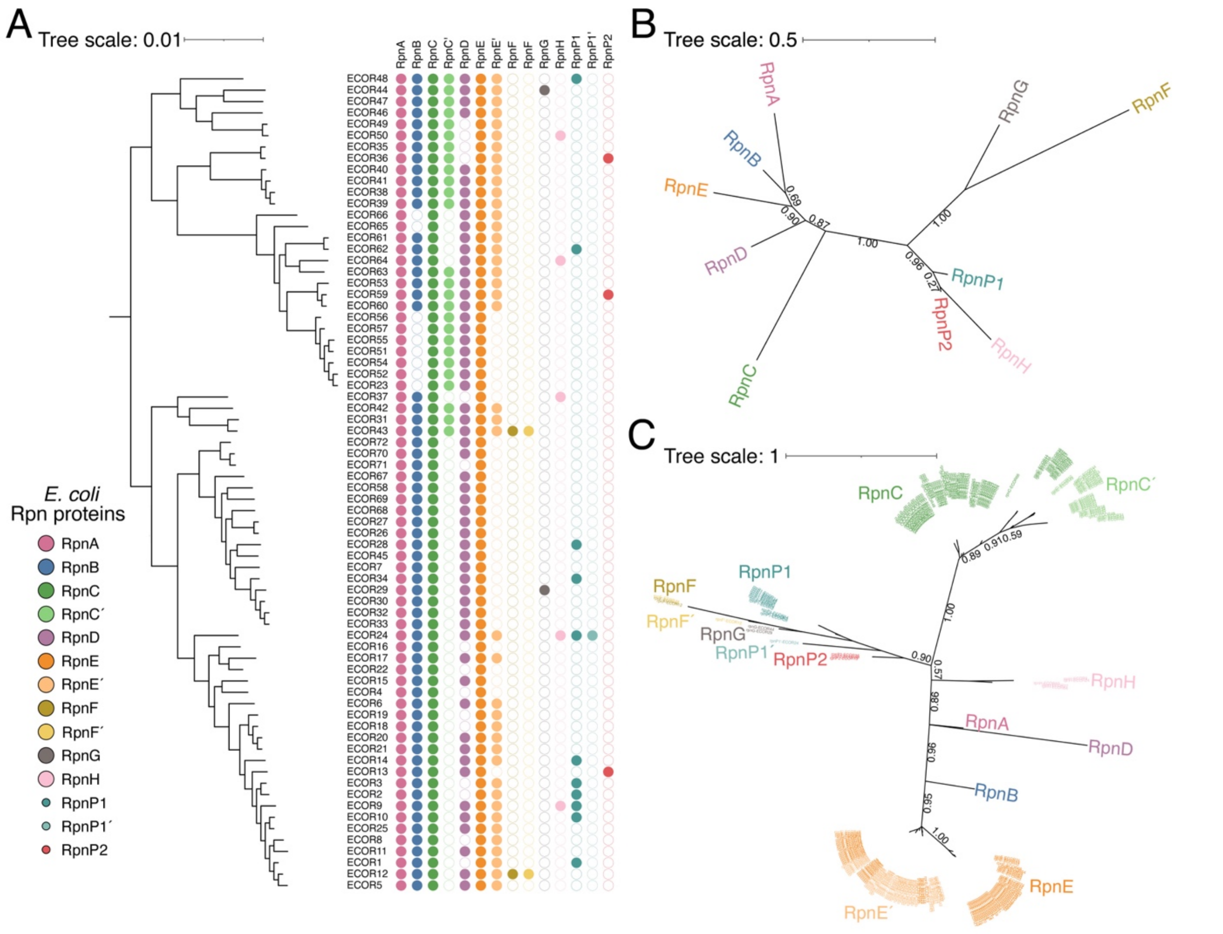

**Fig. S3.** Summary of Rpn proteins found in *E. coli* strains. (*A*) Distribution of Rpn proteins found in the 72 *Escherichia coli* strains of the ECOR (19) collection. RpnA-E are proteins encoded on the chromosome of *E. coli* K-12. RpnF-H also are chromosomally encoded, while RpnP1 and RpnP2 are encoded on plasmids. (*B*) Phylogenetic tree based on the protein sequences of the N-terminal domain of *rpn*$_L$ in *E coli*. (*C*) Phylogenetic tree based on the protein sequences of *rpn*$_S$ and *rpn*$_{S'}$ in *E coli*. For (*B*) and (*C*), the numbers on the Tree branches corresponds to the bootstrap value, which measures the reliability of the topology of the branch. The value should be between 0 and 1.

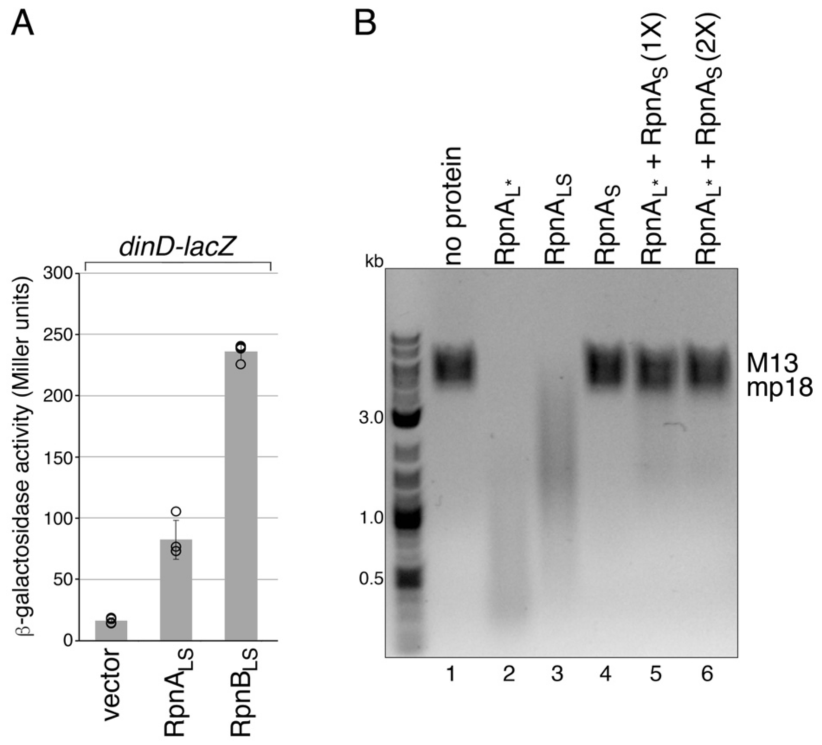

**Fig. S4.** $Rpn_L$ activity requires *C*-terminal domain and active site and is blocked by $Rpn_S$. (*A*) WT $RpnA_{LS}$ and $RpnB_{LS}$ induce *dinD-lacZ* activity. All data represent four independent biological repeats. The ER2170 strain background is WT for the *rpn* genes. As described in Materials and Methods, growth conditions differed somewhat from those in Fig. 3*D* and 3*E*. (*B*) Purified $RpnA_S$ blocks ssDNA endonuclease activity of $RpnA_{L*}$. Purified $RpnA_{L*}$ (lane 2), $RpnA_{LS}$ (lane 3), or $RpnA_S$ (lane 4) or $RpnA_{L*}$ mixed with $RpnA_S$ (lanes 5 and 6) were incubated with M13mp18 ssDNA. Image is inverted from the original. The nuclease assay was repeated at least twice.

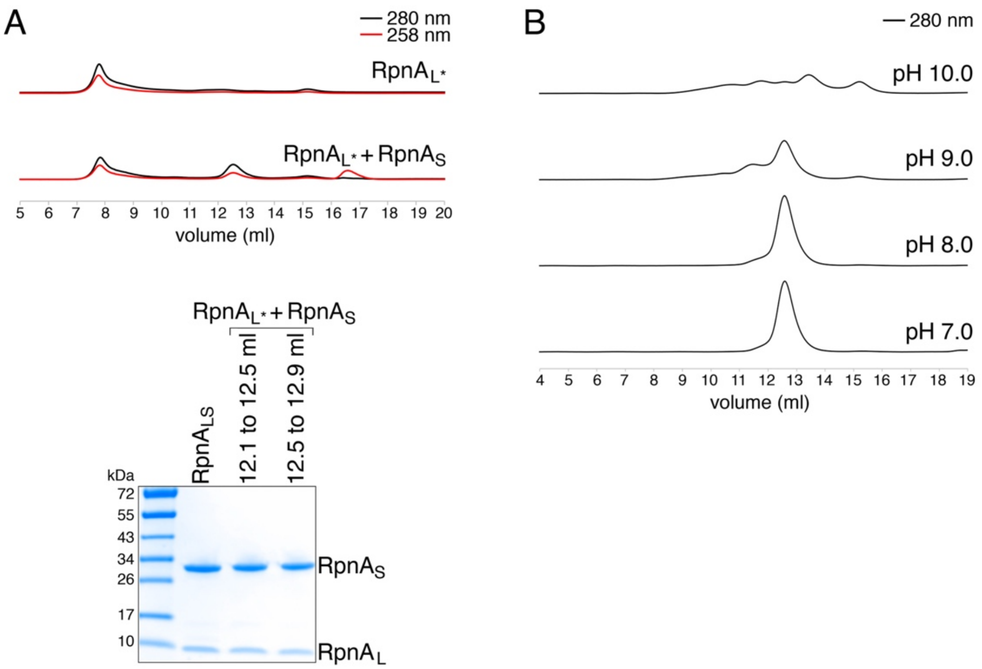

**Fig. S5.** Size exclusion chromatography (SEC) analysis of RpnA$_{L*}$, RpnA $_{L*}$-RpnA$_S$ complex and RpnA$_{LS}$. (*A*) Sizing column of RpnA$_{L*}$ and RpnA$_{L*}$ + RpnA$_S$. The separation was achieved by Superdex 200 Increase 10/300 GL column. RpnA$_{L*}$ + RpnA$_S$ peak fractions corresponding to 12.1 to 12.5 mL and 12.5 to 12.9 mL were pooled and analyzed on a Coomassie-stained Tris-glycine SDS-PAGE gel. (*B*) Effect of pH on oligomerization. RpnA$_{LS}$ was incubated at pH 7.0, 8.0, 9.0, 10.0 at room temperature overnight before analysis by a Superdex 200 Increase 10/300 GL column.

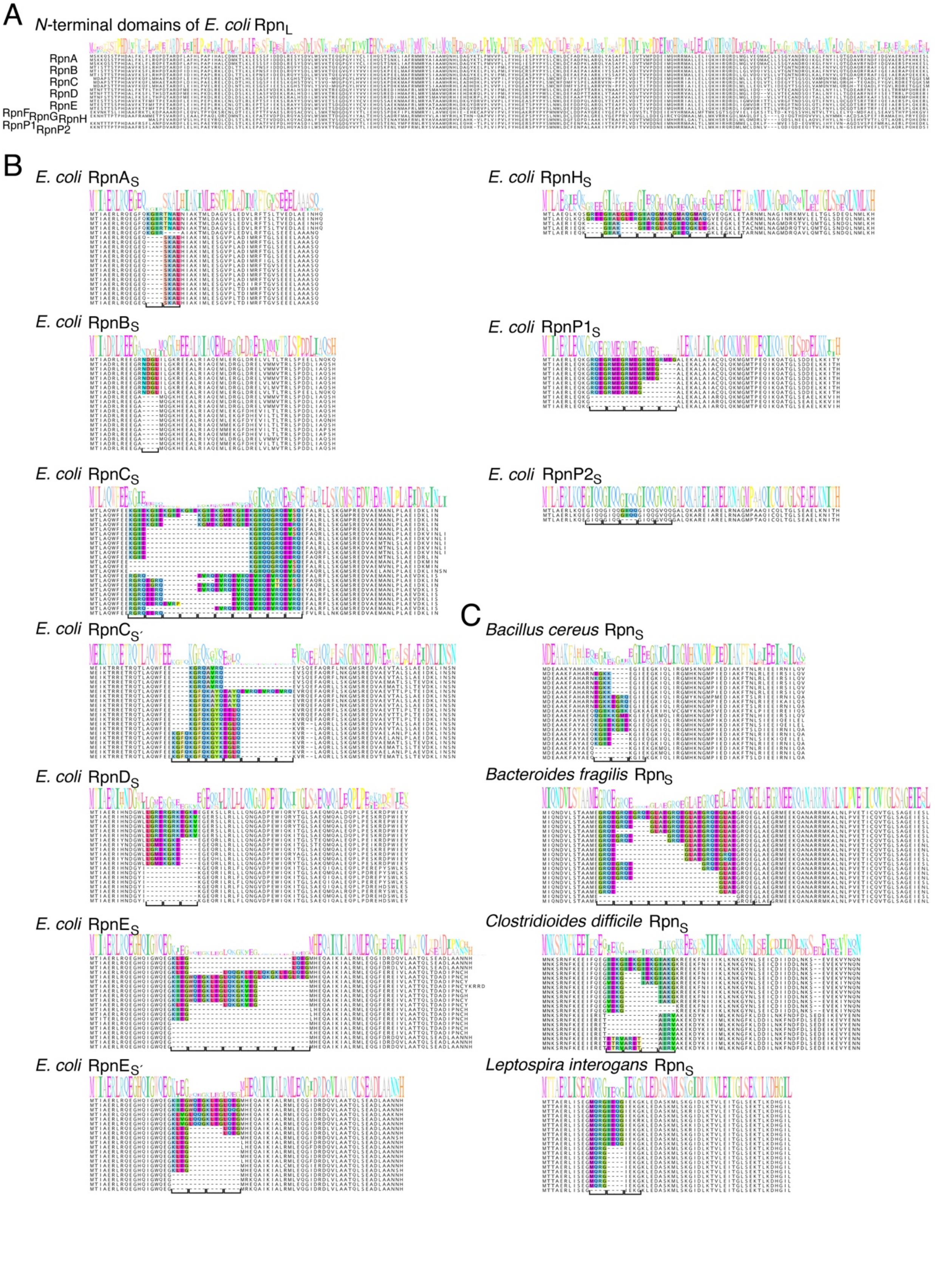
A
N-terminal domains of E. coli RpnL
RpnA
RpnB
RpnC
RpnD
RpnE
RpnF
RpnG
RpnH
RpnP1
RpnP2
B
E. coli RpnAS
E. coli RpnBS
E. coli RpnCS
E. coli RpnCS′
E. coli RpnDS
E. coli RpnES
E. coli RpnES′
E. coli RpnHS
E. coli RpnP1S
E. coli RpnP2S
C
Bacillus cereus RpnS
Bacteroides fragilis RpnS
Clostridioides difficile RpnS
Leptospira interogans RpnS

**Fig. S6.** Alignments of Rpn *N*- and *C*-termini. (*A*) Sequence alignment of *N*-terminal domains of representative $RpnA_L$, $RpnB_L$, $RpnC_L$, $RpnD_L$ and $RpnE_L$ proteins (three each) as well as $RpnF_L$, $RpnG_L$, $RpnH_L$, $RpnP1_L$ and $RpnP2_L$ from *E. coli*. (*B*) Sequence alignment of $Rpn_S$ proteins from *E. coli*. (*C*) Sequence alignment of $Rpn_S$ proteins from indicated Gram+ bacteria. For (*B*) and (*C*), the sequence logo was built based on all the sequences for each group, though only 20 representative sequences are shown for those where more than 20 are available. The four amino acid repeat that varies between strains is colored in the alignment, with a bracket indicating the number of repeats. The alignments were manually curated.

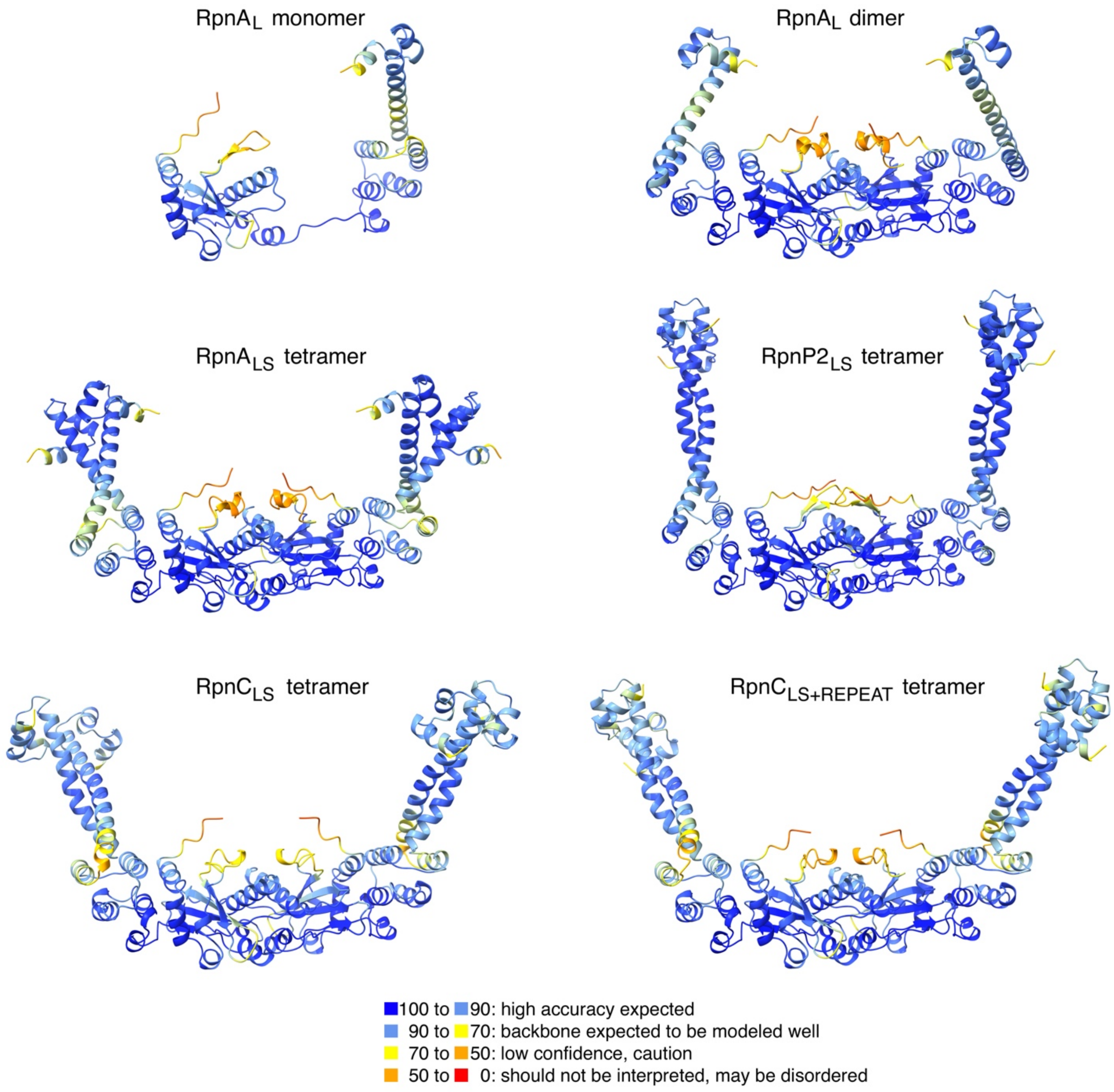

**Fig. S7.** AlphaFold predictions. The AlphaFold predictions for RpnA$_L$ as a monomer and dimer as well as a tetramer with RpnA$_S$ are shown along with the predictions for the RpnP2$_{LS}$, RpnC$_{LS}$ and RpnC$_{LS+REPEAT}$. Key for confidence in predictions is given at the bottom.

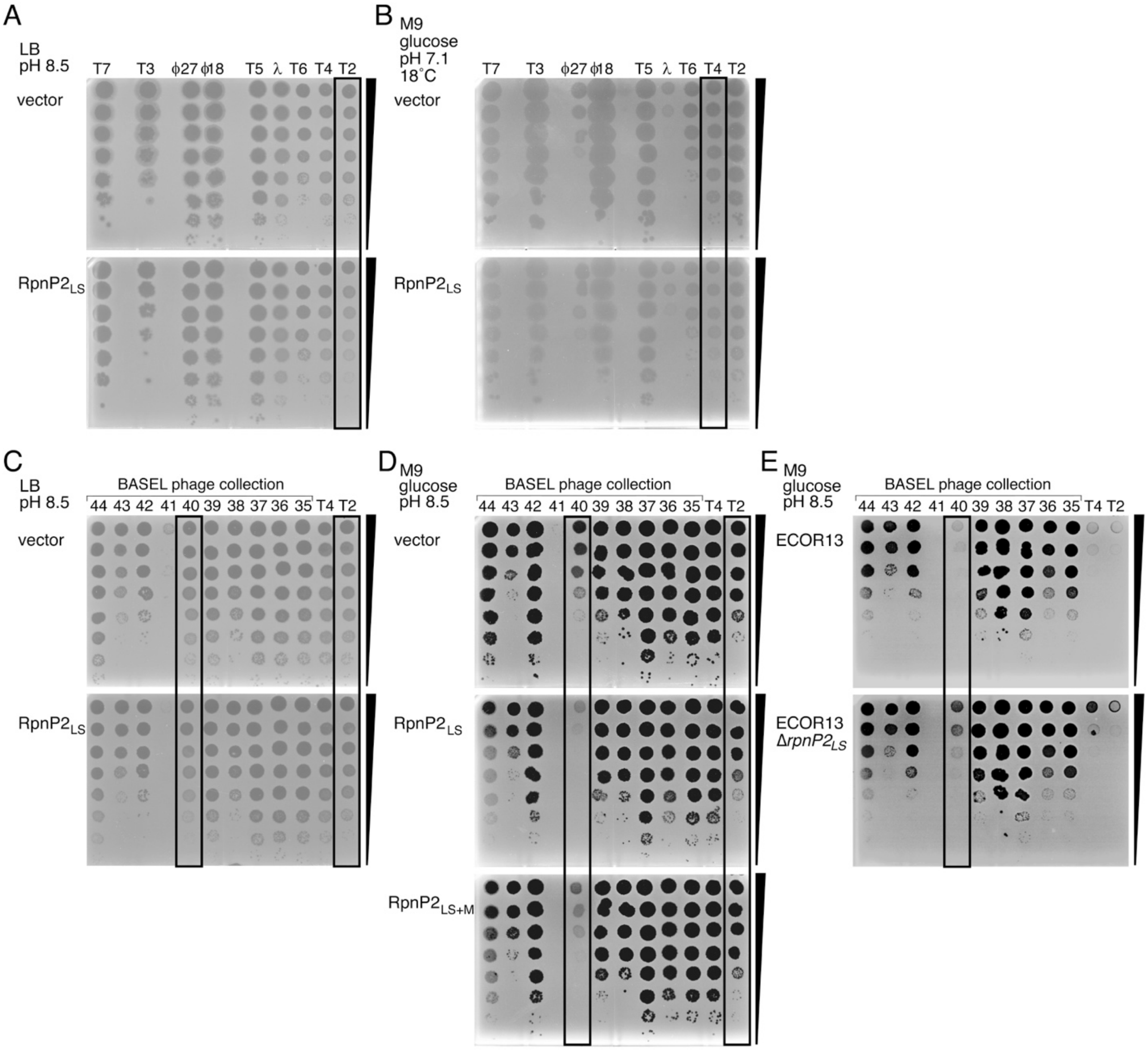

**Fig. S8.** Plaque assays. (*A*) Plaquing of 10-fold serial dilutions of indicated phage on *E. coli* MG1655 cells harboring vector or $RpnP2_{LS}$ grown on LB medium buffered to pH 8.5. (*B*) Plaquing of dilutions of indicated phage on *E. coli* MG1655 cells harboring vector or $RpnP2_{LS}$ grown at 18˚C on M9 glucose medium pH 7.1. (*C*) Plaquing of dilutions of indicated T-even phages on *E. coli* MG1655 cells harboring vector or $RpnP2_{LS}$ grown on LB medium buffered to pH 8.5. (*D*) Plaquing of dilutions of indicated T-even phages on *E. coli* MG1655 cells harboring vector, RpnP2$_{LS}$ or RpnP2$_{LS+M}$ carrying D14A, R66A, E86A, and Q88K mutations on M9 glucose medium buffered to pH 8.5. While plaque size for this mutant is comparable to WT for T2 and BASEL phage #37 and 38, the mutant strain retains some resistance to BASEL phage #40. (*E*) Plaquing of dilutions of indicated T-even phages on *E. coli* ECOR13 or ECOR13 Δ*rpnP2*::*cm* cells grown on M9 glucose medium buffered to pH 8.5. A representative image of at least two independent biological repeats is shown. The black wedge denotes the 10-fold dilution series.
